# Cell surface fluctuations regulate early embryonic lineage sorting

**DOI:** 10.1101/2020.08.16.250084

**Authors:** Ayaka Yanagida, Christopher Revell, Giuliano G. Stirparo, Elena Corujo-Simon, Irene M. Aspalter, Ruby Peters, Henry De Belly, Davide A. D. Cassani, Sarra Achouri, Raphael Blumenfeld, Kristian Franze, Ewa K. Paluch, Jennifer Nichols, Kevin J. Chalut

## Abstract

In development, lineage segregation of multiple lineages must be coordinated in time and space. An important example is the mammalian inner cell mass (ICM), in which the primitive endoderm (PrE, founder of the yolk sac) physically segregates from the epiblast (EPI, founder of the foetus). The physical mechanisms that determine this spatial segregation between EPI and PrE are still poorly understood. Here, we identify an asymmetry in cell-cell affinity, a mechanical property thought to play a significant role in tissue sorting in other systems, between EPI and PrE precursors (pEPI and pPrE). However, a computational model of cell sorting indicated that these differences alone appeared insufficient to explain the spatial segregation. We also observed significantly greater surface fluctuations in pPrE compared to pEPI. Including the enhanced surface fluctuation in pPrE in our simulation led to robust cell sorting. We identify phospho-ERM regulated membrane tension as an important mediator of the increased surface fluctuations in pPrE. Using aggregates of engineered cell lines with different surface fluctuation levels cells with higher surface fluctuations were consistently excluded to the outside of the aggregate. These cells behaved similarly when incorporated in the embryo. Surface fluctuations-driven segregation is reminiscent of activity-induced phase separation, a sorting phenomenon in colloidal physics. Together, our experiments and model identify dynamic cell surface fluctuations, in addition to static mechanical properties, as a key factor for orchestrating the correct spatial positioning of the founder embryonic lineages.

## Main

An essential event in the development of a mammal is the segregation of the EPI, which will form the foetus, from the extraembryonic tissues that manage implantation, nutrition and patterning of the foetus. The first step of this process is the formation of the blastocyst, which has been well-described in mouse (1-3). The blastocyst forms as the outside cells of the preimplantation embryo differentiate into trophectoderm (the source of the placenta) and cavitation occurs (Fig. 1a). At this point, the inside cells comprising the inner cell mass (ICM), are aggregated and firmly adhered to the trophectoderm on the proximal pole of the blastocyst (Fig. 1a, E3.5). Subsequently, a subpopulation of ICM cells become sensitive to FGF4, heralding PrE bias (4, 5). Within the uterus, and ex vivo, precursors of the EPI and PrE emerge in a spatially random manner (6-8). Coincident with identity acquisition, the cells physically sort, resulting in PrE establishing a single layer of cells covering the cavity-facing surface of the ICM with the EPI enclosed between the PrE and polar trophectoderm (8). The chemical signalling requirements for fate specification are well-understood: FGF4-dependent ERK activation is necessary and sufficient for PrE specification in the mouse (9). Less is known about how proper positioning of PrE is achieved. It is known that once all PrE cells are on the cavity-facing surface of the ICM, they undergo an aPKC-dependent epithelialization that maintains the positioning (10). However, how PrE cells mechanically segregate from the EPI in the first place is currently not understood (Fig. 1a).

**Fig. 1.**
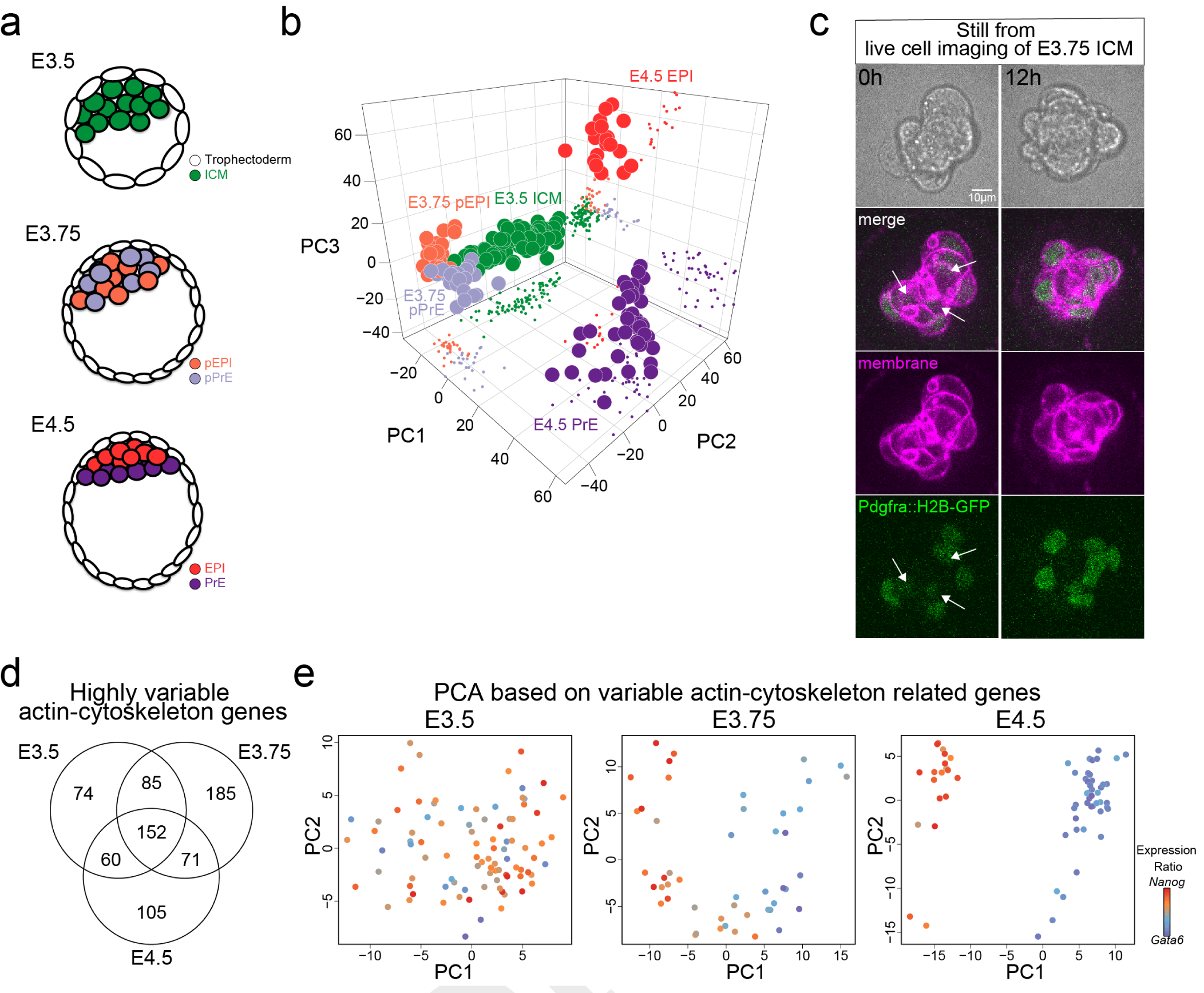
EPI and PrE begin to segregate at E3.75. (a) Schematic of EPI and PrE segregation in blastocysts. (b) Principal Component Analysis (PCA) plot of E3.5, E3.75 and E4.5 cells coloured according to their stage. Each dot represents a single cell. (c) Images of an isolated E3.75 *Pdgfra*^*H2B-GFP/+*^*mTmG*^*+/-*^ ICM cultured *ex vivo*, taken as stills from movies (Supplementary movie 1). T = 0h and 12h show the EPI and PrE sorting stage and completed sorting stage respectively. *Pdgfra*^*H2B-GFP*^ was expressed in PrE nuclei (green) and mTmG at the cell membrane (magenta). Arrows indicate pPrE cells located inside the ICM. (d) Venn diagram of the number of highly variable actin-cytoskeletal genes in E3.5, E3.75 and E4.5 ICM cells. (e) PCA plot of E3.5, E3.75 and E4.5 ICM cells based on the highly variable cytoskeletal genes (E3.5: n = 371, E3.75: n = 493, E4.5: n = 388, log_2_ FPKM > 1, logCV2 > 0.5) coloured according to the ratio of *Nanog* to *Gata6* expression. Each dot represents a single cell of ICM cells.

Several mechanical mechanisms of sorting have been proposed previously, including differential adhesion (11) and differential surface tension (12), and differential cell-cell affinity (13-16). Here, we examined all these possibilities in the context of sorting in the ICM, and found that all of these static cell mechanical properties appeared insufficient to explain robust segregation of the PrE lineage from the EPI lineage. Instead, through a combination of experiments and physical modelling, we uncovered that enhanced surface fluctuations, an intrinsically dynamic cellular mechanical property, in the PrE lineage are an essential factor in facilitating the segregation of these early embryonic lineages.

To determine the most relevant stage to investigate ICM sorting, we used RNA sequencing to analyse the gene expression of single ICM cells at E3.75, and combined it with previous analyses performed at E3.5 and E4.5 (17). Principal component analysis (PCA) revealed stage-specific clusters, indicating that in the E3.75 ICM, progenitors with specific embryo lineages, pEPI and pPrE, that are just beginning to become distinct (Fig. 1b and Fig. S1a-c).

To identify a simplified system to study this lineage segregation, we first confirmed, as shown in (18), that pEPI and pPrE cells in ICMs isolated from E3.5 or E3.75 blastocysts can segregate and commit to EPI and PrE in culture without trophectoderm (Fig. S2a). The majority of E3.5 ICM formed ‘miniblastocysts’ (Fig. S2b) containing cavities, with some external cells expressing the trophectoderm marker, CDX2 (Fig. S2c). The later E3.75 ICMs formed embryoid body-like structures with no cavity or CDX2 expressing cells. After one day in culture, the PrE enveloped the EPI, confirmed using immunofluorescence (Fig. S2d, e). Taken together, our data confirm proper fate segregation and maturation in isolated E3.75 ICMs in the absence of trophectoderm.

To study the dynamics of segregation of pEPI and pPrE cells in E3.75 ICMs, we generated time-lapse movies of isolated ICMs from embryos expressing both a PrE lineage reporter, *Pdgfra*^*H2B-GFP*^ (19), and a plasma membrane-localised reporter, mTmG (20) (Fig. 1c and Supplementary movie 1); pPrE cells that were initially randomly distributed sorted to the surface of the ICM.

To understand what drives the observed sorting, we first considered three mechanisms suggested in the literature: differences in cell-cell adhesion, in migration, or in cell polarity (21). Importantly, cell-cell adhesion forces are known to play only a small role in tissue sorting (14). Moreover, E-cadherin, an important regulator of cell-cell adhesion in early development (22), has also been shown not to be differentially distributed at the protein level and to be unnecessary for cell sorting in the blastocyst (23). We also found that E-cadherin is not differentially expressed transcriptionally between pEPI and pPrE at E3.75 (Fig. S3a, showing that N- and P-cadherin are also not differentially expressed). Taken together, we conclude that cell-cell adhesion is unlikely to play more than a minor role in the sorting of the mouse ICM.

A contribution of directed migration to sorting is more difficult to rule out definitively; however, there are scarce indications in images of ICM sorting (8) that pPrE cells in the ICM display elongated shapes typical of mesenchymal migration. Nevertheless, it is possible that they are capable of amoeboid migration in confinement (24), which would be difficult to detect from shape analysis alone. We thus assessed the ability of ICM cells to migrate using a 3-dimensional confinement device (25). We found that there was almost no detectable migration of ICM cells, regardless of the level of confinement (Fig S3b-e), even though other cells types can migrate efficiently in similar conditions (26). This suggests that ICM cells do not have a high level of migration competence, making migration an unlikely candidate to drive robust cell sorting.

Polarity has been shown to be important in establishing the PrE lineage and positioning (10, 21). Polarity molecules, such as Lrp2 and Dab2 (27), are important for instructive signalling in the embryo. However, no direct connection has been shown between differential expression of polarity molecules and mechanical properties that could drive sorting itself. Importantly, aPKC, a polarity protein that has been suggested to be essential for maturation of the blastocyst (10), did not display significant differential expression between pEPI and pPrE at E3.75/E4.0, either at the transcriptional or protein level (Fig. S3f, g). aPKC is primarily cytoplasmic in the ICM up to E4.0, by which time sorting is near completion, while it is clearly surface-localised at E4.5 (Fig. S3g). This is in line with previous work revealing that aPKC is expressed in the ICM only after all PrE cells are localised at the surface (10). Yet, PrE cells stably localise to the surface as individual cells (8), prior to aPKC polarization, suggesting that aPKC is primarily important for reinforcing polarized epithelia (28) rather than the initial sorting of the cells. Thus, how PrE cells become sequestered to the outside of the ICM such that they can form an epithelium remains unresolved (21).

Given that differences in cell-cell adhesion, polarity and migration do not seem to be good candidates to propel robust cell sorting in the blastocyst, we turned to another candidate that has been suggested to drive cell sorting, cell mechanics (29). First, we probed our transcriptomics data for changes in the actin cytoskeleton and its regulators. E3.75 ICM cells, compared to ICM cells at other blastocyst stages, showed the most highly modulated actin cytoskeleton-related genes (Fig. 1d, Supplementary Information Table 1). PCA of each stage based on variable actin cytoskeleton-related genes showed that their expression became distinct at E3.75 (Fig. 1e), coinciding with pEPI and pPrE cells sorting, suggesting that there may indeed be mechanical asymmetries arising between pPrE and pEPI at E3.75.

To probe further into the possibility of a mechanical asymmetry mediating sorting, we first explored differences in surface tension, proposed to be a mechanical driver of cell sorting (12). To explore this possibility in the ICM, we used an atomic force microscope to measure cellular surface tension (30). The surface tension of pEPI and pPrE isolated from E3.75 ICMs was highly variable, but no significant differences were detected (Fig. S4a). Cortical tension is primarily controlled by myosin II activity (31). We thus assessed the levels of phosphorylated myosin regulatory light chain (pMRLC), as MRLC phosphorylation is a key regulator of myosin activity (32), and found no difference between pEPI and pPrE (Fig. S4b). Together, these data suggest that surface tension alone is unlikely to be a major factor in driving pEPI/pPrE sorting.

Another suggested mechanical driver of segregation in developing tissues is differential cell-cell affinity, which is determined by the force balance between cell-cell adhesion, cell surface tensions and interfacial tension at cell contacts (13-16) (Fig. 2a). Physical modelling suggests that in small multicellular systems, differences in cell-cell affinity are sufficient to drive cell sorting (33). Indeed, as we showed previously, differential affinity between two cell types is an excellent predictor of cell sorting (33). Differences in cell-cell affinity can be quantified by differences in contact area or angles (Fig. 2b). Thus, to test whether differential affinity could control ICM sorting, we measured the external contact angles between pEPI and pPrE doublets made by aggregating two E3.75 *Pdgfra*^*H2B-GFP*^ ICM cells (Fig. 2b, c). The external contact angles of homotypic pEPI cell doublets (pEPI::pEPI) were significantly larger than those of both pPrE::pPrE and pEPI::pPrE (Fig. 2d). An affinity parameter, *β*, between two cell types can be calculated based on these angles (Fig. 2a, Supplementary information). The affinity parameter reflected the cell-cell affinity differential between different types of doublets and was found to be:

**Fig. 2.**
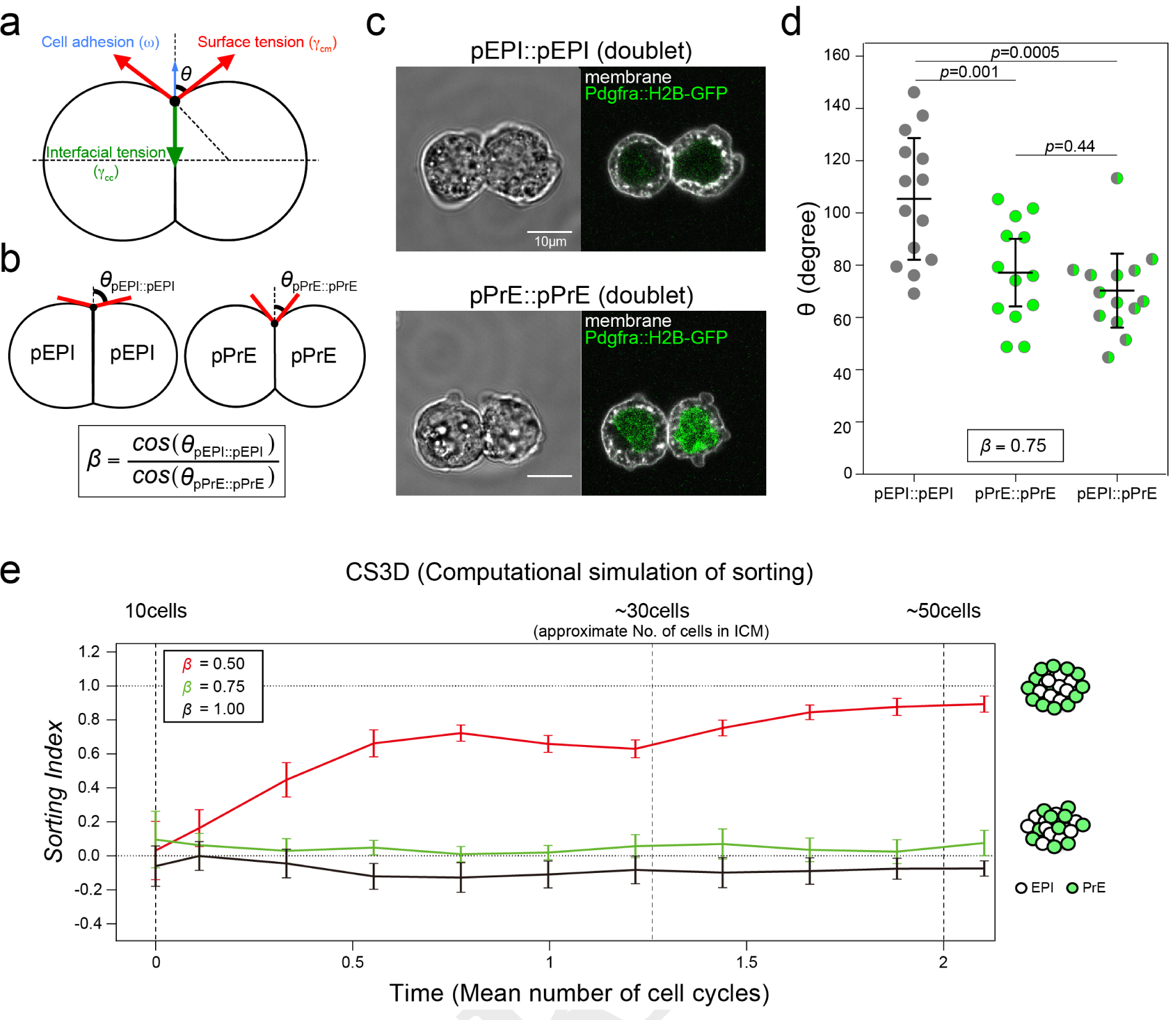
Cell-cell affinity of pEPI is higher than pPrE, but insufficient for lineage segregation. (a) Schematic of two ICM cells forming a doublet. The shape of the doublet is determined by the balance of surface tensions *γ*_*cm*_ at the cell-medium interface, interfacial tension *γ*_*cc*_ at the cell-cell interface, and cell-cell adhesion force *ω*. (b) Schematic of pEPI or pPrE cells homotypic doublet showing how *β* is used as a measure for cell-cell affinity. *θ*_*pEP I*::*pEP I*_ shows the pEPI::pEPI external contact angle, and *θ*_*pP rE*::*pP rE*_ shows the PrE::pPrE external contact angle. (c) Representative images of a pEPI (pEPI::pEPI) and a pPrE (pPrE::pPrE) homotypic doublet, formed by dissociated single cells from E3.75 *Pdgfra*^*H2B-GFP/+*^ blastocysts. pPrE expressed *Pdgfra*^*H2B-GFP*^ at nuclei (green). Plasma membrane labelled with a membrane dye, CellMask Deep Red (false-coloured white). (d) *θ* of the different types of doublets that can be formed from E3.75 pEPI and pPrE cells. Each dot represents the mean of both sides of external contact angles. The data is combined over N = 3 independent experiments, and *β* as calculated from the mean *θ*_*pEP I*::*pEP I*_ and *θ*_*pP rE*::*pP rE*_ as 0.75 from equation in (b). P-values calculated by 2-way ANOVA using cell type and replicate number (N = 3) as variables. (e) 3D force-based cell sorting simulation (CS3D) simulation of EPI and PrE sorting applied with differential affinity ratio *β, β* = 1.0 indicating no difference in affinity between pEPI and pPrE assuming a system evolving from 10 to 50 cells, which represents slightly more than two cell divisions. *Sorting index = 0*.*0* indicates random sorting and *Sorting index = 1*.*0* indicates complete sorting with PrE located on the outside.

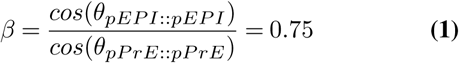

*β <* 1 indicates that the affinity is greater in pEPI::pEPI than pPrE::pPrE, and that pPrE should thus sort to the outside of an aggregate.

To test whether the measured affinity parameter is sufficient to account for the segregation of pEPI and pPrE cells, we used a 3D computational model based on the subcellular element method, termed 3D force-based cell sorting simulation (CS3D, described in (33) and also in Supplementary Information). Briefly, each individual cell is modelled as a group of infinitesimal elements, interacting via nearest-neighbour forces. CS3D also incorporates cell growth and division, and enables a multiscale modelling of inter- and intra-cellular interactions in multicellular aggregates such as the ICM. To score sorting in the aggregates, we established a sorting index ((33) and Extended Supplementary Methods). The sorting index has a range of -1 to 1, where -1 indicates pEPI cells on the outside, 0 indicates random cell positioning (no sorting), and 1 indicates pPrE cells on the outside. Using CS3D, we simulated sorting in the ICM with our experimentally observed value of *β* = 0.75, with cells dividing from 10 to up to 50 cells. These numbers represent the approximate beginning and end number of cells in the ICM between E3.5 and E4.5, while the average ICM at E3.75 possesses 30 cells (34). Surprisingly, in our simulations, the measured level of differential cell-cell affinity was not sufficient to lead to pEPI and pPrE sorting, and a much stronger differential affinity (*β <* 0.5) was required to efficiently sort cells within the experimental timeframe (Fig. 2e). Our model thus implied that the experimentally measured affinity differential is not sufficient to support robust sorting of pEPI and pPrE. Taken together with all of the other results, this suggested we were missing a key parameter.

Interestingly, consistent with earlier reports from the late blastocyst (35), we noticed that mid blastocyst pPrE cells displayed a less smooth morphology compared to pEPI cells (Fig. 3a, Fig. S5). We thus asked whether these differences were due to increased dynamic cell surface fluctuations in pPrE. To answer this, we isolated pEPI and pPrE cells from several stages of early ICM, including E3.75, in a *mTmG*^*+/-*^ *Pdgfra*^*H2B-GFP*^ background, and live-imaged them for 5 minutes. We then quantified the amplitude of surface fluctuations of the cells (Fig. 3b, c and Supplementary Methods). Single pPrE cells exhibited significantly larger surface fluctuations than pEPI (Fig. 3c). Notably, this quantification does not distinguish between different types of cellular protrusions; but we observed that the primary manifestation of the surface fluctuations was blebbing (Supplementary movie 2, 3, Fig. S5).

**Fig. 3.**
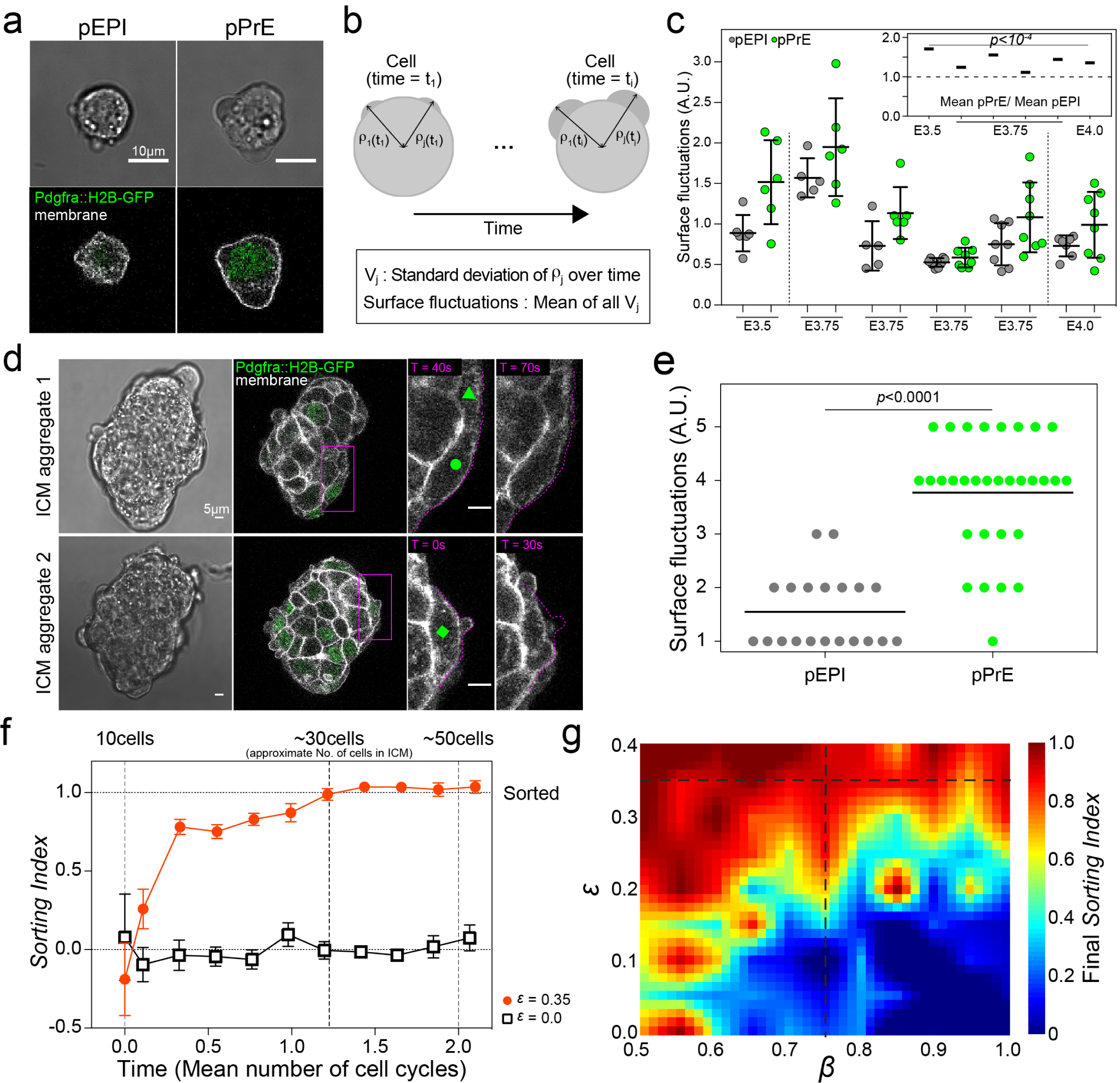
Higher surface fluctuations in pPrE regulate ICM sorting. (a) Representative images of dissociated E3.75 pEPI and pPrE cells from *Pdgfra*^*H2B-GFP/+*^ *mTmG*^*+/-*^ mice, showing that pPrE typically exhibit more blebbing than pEPI. As opposed to E4.5 PrE, the blebs tend to be large blebs. pPrE expressed *Pdgfra*^*H2B-GFP*^ at nuclei (green). Plasma membrane labelled with membrane dye, CellMask Deep Red (white). (b) Schematic showing how surface fluctuations are calculated. (c) Single E3.5, E3.75 and E4.0 pEPI and pPrE surface fluctuations. The amplitude of surface fluctuations was calculated using images every ten seconds over a total of five minutes and normalised by the total mean of ICM cell surface fluctuations over all time points and conditions. The inset presents only means of each experiment, showing a consistently higher level of surface fluctuations in pPrE through all states of ICM sorting. P-value calculated by 2-way ANOVA using cell type and replicate number as variables, p-value reported in inset is for cell type (PrE versus EPI lineage). (d) Representative images of isolated ICM aggregations, each aggregation comprising 3 ICMs, from E3.75 *Pdgfra*^*H2B-GFP/+*^ *mTmG*^*+/-*^ embryos, taken as stills from movies (Supplementary movie 2-3). Each outside cell was ranked single-blind from 1 to 5, 5 indicating a high level of surface fluctuations and 1 indicating no visible surface fluctuations. Cells with green triangle, circle, and square indicating surface fluctuations = 2, = 3 and = 5 respectively. (e) Blind rank analysis of surface fluctuations of pPrE and pEPI cells. P-value calculated by 1-way ANOVA. (f) Time series plot of CS3D simulation from 10 to 50 cells, assuming symmetric division, using the experimentally measured value of *β* = 0.75. *ϵ* = 0 means no difference in surface fluctuations between pEPI and pPrE. *ϵ* = 0.35 is the measured ratio of surface fluctuations of pPrE to pEPI. Each parameter set is averaged over N = 4 runs. The horizontal dotted line (*Sorting Index = 1*.*0*) shows the threshold beyond which sorting is complete, with pPrE on the outside. (g) Phase spaces of the final sorting index for 30 cells’ aggregates in *ϵ* and *β* space, with a resolution of 0.05 on each axis. The dotted lines (*ϵ* = 0.35, *β* = 0.75) represent the experimentally measured parameters.

To examine whether differences in surface fluctuations were observable in the multicellular context, we monitored cellular membrane dynamics in ICMs isolated from blastocysts. Surface fluctuations were clearly visible on the outer edge of cell aggregates. In order to ensure a sufficient number of cells located on the outside layer to perform a quantification of surface fluctuations, we aggregated three isolated ICMs from E3.75 *mTmG*^*+/-*^ *Pdgfra*^*H2B-GFP*^ blastocysts (Fig. 3d). We then performed a blinded quantification of surface fluctuations in the aggregates, and found that pPrE cells on the outer layer of the ICM aggregate exhibited significantly higher surface fluctuations than pEPI cells similarly located (Fig. 3e and Supplementary movie 4).

Upon consideration of our data, we speculated that differences in surface fluctuations could contribute to segregating pEPI and pPrE cells. To test this possibility, we extended the CS3D simulations to incorporate random cell surface fluctuations. We established a dimensionless parameter *ϵ* which represents the ratio of surface fluctuations in pPrE to pEPI (Extended Supplementary Information). We used the measured surface fluctuations *ϵ* of pEPI and pPrE to establish an estimate for *ϵ*, finding that *ϵ* ≈ 0.35 from the data (Supplementary Materials). We simulated pEPI and pPrE cell sorting from 10 to 50 cells using CS3D with *β* = 0.75 and *ϵ* = 0, corresponding to equal surface fluctuations in PrE and EPI cells, or the experimentally observed fluctuations differential *ϵ* = 0.35. No sorting was observed for *ϵ* = 0. However, we observed thorough and robust sorting for *ϵ* = 0.35 (Fig. 3f). We then assembled a phase space of the sorting index for the range *β* = [0.5, 1.00] and *ϵ* = [0.0, 0.40] to cover a wide range of experimental parameters. It is clear from the phase space that, though we see moderate segregation without a fluctuation differential as *β* approaches sufficiently extreme values of 0.5, the cells are capable of sorting even if *β* = 1 as long as the pPrE cells have significantly larger surface fluctuations (Fig. 3g). Thus, our model suggests that a differential in surface fluctuations could strongly contribute to the sorting of pEPI and pPrE.

In order to direct our hypothesis that increased surface fluctuations lead to sorting *in vivo*, we sought a candidate to increase fluctuations that did not significantly affect cortical tension. For this, we considered the effective membrane tension, which is primarily regulated by the level of attachment between the plasma membrane and the underlying cortex (36), with the Ezrin-Radixin-Moesin (ERM) protein family (37) playing a key role. Thus, we focused on ERM, exploiting the fact that depleting ERM decreases effective membrane tension (30, 37), leading to enhanced blebbing, while having little effect on cortical tension (38). We thus performed a triple knockdown of ERM in mouse embryonic stem (ES) cells, and confirmed that it led to higher levels of surface fluctuations (Fig. 4a). We then used chimaeras in which either negative control (NC) siRNA treated ES cells or ERM-depleted ES cells were injected to E3.5 blastocysts to determine whether changing the levels of surface fluctuations would affect epiblast incorporation. For this, we assessed the cultured blastocyst at E4.0, when injected ES cells have been shown to populate the epiblast (39). Indeed, wild type ES cells were primarily situated in the epiblast. In contrast, we found that a significant fraction of the ERM knockdown cells were incorporated into the ICM, but in the outer, putative PrE, layer (Fig. 4b, c). This suggests that ES cells with enhanced surface fluctuations could not properly incorporate into the epiblast. This strongly supports the hypothesis that surface fluctuations play an important role in sequestering cells to the outer ICM layer.

**Fig. 4.**
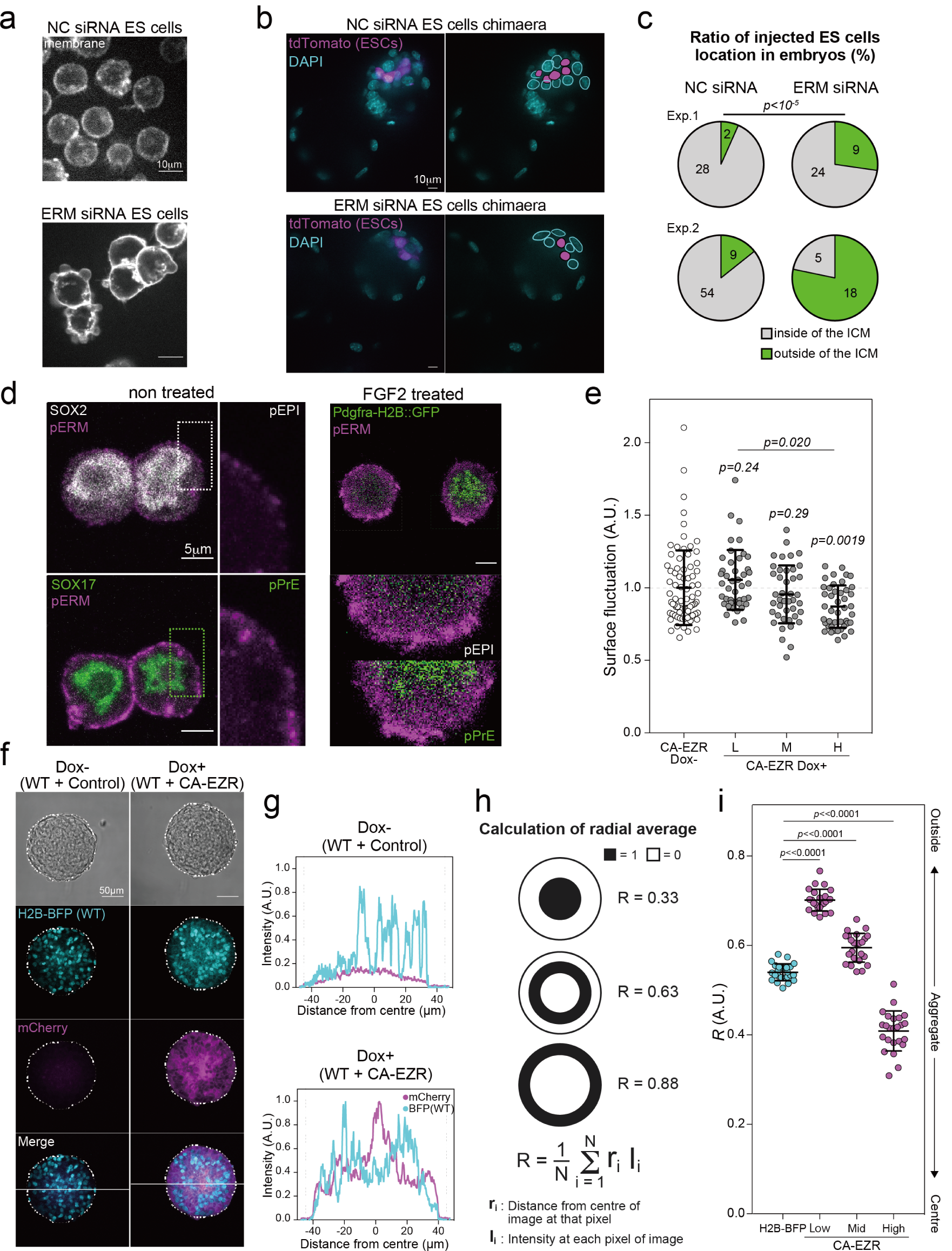
ERM activity mediates surface fluctuations and cell segregation in vivo and in aggregates. (a) Representative images of negative control (NC) siRNA treated ES cells, and ezrin, radixin and moesin (EZR) siRNA treated ES cells. (b) Representative images of NC siRNA-treated tdTomato ES cells chimaera, and ERM siRNA treated tdTomato ES cells chimaera, both at E4.0. Nuclei were stained with DAPI (cyan). ES cells were labelled with tdTomato (magenta). (c) The pie charts of injected NC siRNA or ERM siRNA treated ES cells location in the ICM of the chimaera blastocysts. For further details about number of embryos, live cells, etc, see *Supplementary Table 4*. The number of ES cells located at the outside (green) or the inside (grey) of ICM was counted using the fixed chimaera blastocyst images. P-value calculated using Fisher’s exact test. (d) Representative images of pERM expression in pEPI and pPrE (magenta) with or without FGF2. pEPI expressed SOX2 (white). pPrE expressed SOX17 or *Pdgfra*^*H2B-GFP*^ at nuclei (green). pERM is clearly more highly expressed, and highly variable at the surface, in pPrE. (e) Surface fluctuations of single cells, N = 2 experiments, of CA-EZR ES cells with or without Dox in 2i+LIF. L, M, and H mean low, medium and high expression of mCherry as assessed by the 3-quantiles of expression in the mCherry expressing cells. Surface fluctuations were normalized by the mean of the Dox-surface fluctuations in each of the N = 2 experiments. P-values were calculated using 1-way ANOVA, with the p-values above each group representing the outcome of pairwise comparison with Dox-. (f) Representative images of CA-EZR ES cells and WT H2B-BFP ES cells aggregated with or without Dox. ES cell aggregates were cultured on low attachment dishes in N2B27 with 2i+LIF for one day. (g) Representative comparison of BFP and mCherry signals in the CA-EZR and H2B-BFP ES cells aggregates with or without Dox, using the line across the images in (f). (h) Schematic showing how the dipole moment R is calculated, along with model examples of R for distributions shown. (i) Dipole moment R of aggregates of CA-EZR and H2B-BFP ES cells. P-value was calculated by pairwise comparison using 1-way ANOVA (N = 24 aggregates).

We next asked why there might be enhanced surface fluctuations in pPrE. As the fluctuations manifested mostly as blebs, the most likely candidates are increased cortical tension or decreased membrane tension (40). As cortical tension does not appear to be significantly different between EPI and PrE lineages (Fig. S4), overall differences in cortical tension are unlikely to play a prominent role. Thus, we turned to effective membrane tension. Surprisingly, we found effective membrane tension to be higher in pPrE (Fig. S6a). Correspondingly, we found that there were much higher overall levels of phosphorylated ERM (pERM), the active form of ERMs, in pPrE than pEPI cells (Fig. 4d). Interestingly, we also found that application of exogenous FGF, which is the primary regulator of fate specification in the ICM (9), significantly increased ERM activity and cell surface fluctuations across all cells (Fig. 4d Fig. S7 and Supplementary movies 5, 6). These observations appeared counter-intuitive at first, as high membrane tension generally limits blebbing (40). Significantly, however, along with the higher pERM levels, we observed that there was also a high degree of spatial variability in pERM levels along the cell boundary in pPrE cells (Fig. 4d). High pERM spatial heterogeneity was also observed in the E3.75 blastocyst (Fig. S6b-e). Furthermore, we found that the temporal variability in membrane tension at the single-cell level correlates with the amount of blebbing in the cell (Fig. S6f, g), and observed a higher variation in the thickness of the actin cortex of pPrE cells (Fig. S8). Importantly, heterogeneities in ERM levels and in cortex organization can both promote blebbing (40). Therefore, we speculate that variability in membrane tension, and possibly in cortex organization, is connected to the increased blebbing in the PrE lineage.

To further test the hypothesis that ERM-regulated cell surface fluctuations control cell sorting, we used a Dox-inducible constitutively active Ezrin (EzrinT567D-IRES-mCherry, or CA-EZR for short) mouse ES cell line. We found that, upon Dox induction, cellular mCherry levels anticorrelated with the variability of pERM in the membrane (Fig. S9a, b), indicating that the higher the levels of CA-EZR, the less variable the pERM in the membrane. Concomitantly, we found that a higher expression of mCherry corresponded to lower surface fluctuations (Fig. 4e). We exploited the fact that the CA-EZR cells provide a system possessing a range of surface fluctuations (Fig. S9a, b) and performed cell aggregation assays to directly assess how cell surface fluctuations affect sorting. As a control, we used an H2B-BEP ES cell line that displays lower levels of cell surface fluctuations compared to the CA-EZR line (Fig. S9c). We mixed the control ES cells with the CA-EZR ES cells at a 1:1 ratio and cultured for one day with or without Dox (Fig. 4g). We quantified sorting by calculating the normalised average distance of the mCherry signal from the centre of the aggregate, R (Fig. 4h, i). Using this measure and thresholding to determine low-, mid-, and high-expressing CA-EZR cells, we found that low-expressing CA-EZR cells, which have enhanced surface fluctuations compared to controls, were preferentially found on the outside of the aggregate, while high expressing CA-EZR cells, which have reduced surface fluctuations compared to controls, localised to the inside of the aggregate (Fig. 4j). Taken together, our data strongly support our hypothesis that differences in surface fluctuations lead to cell sorting, with cells possessing larger surface fluctuations sorting to the outside of an aggregate.

There has been a growing consensus that spatial segregation of embryonic cell lineages is typically driven by affinity asymmetries at cellular interfaces (13-15, 30, 41), though a recent report found spatial segregation with no clear affinity asymmetries (42). We show here that, although there is affinity asymmetry between emergent EPI and PrE lineages, these asymmetries appear to be insufficient to explain their spatial segregation. Indeed, our simulations suggest that for cell sorting to occur within a timeframe relevant to development, a high degree of affinity asymmetry is required, which is not necessarily achieved for all embryonic tissues, including EPI and PrE in the mouse blastocyst. We cannot exclude the possibility that the small differential cell-cell affinity leads to an ‘outside bias’ for the PrE lineage, but it does not explain why PrE cells remain there long enough to form the later-emerging polarized epithelium.

This is the first time, to our knowledge, that a mechanism to explain spatial segregation in tissue is based not on differences in static physical properties, but instead on dynamic mechanical fluctuations, such as those introduced by cellular blebs. The mechanism we propose here is reminiscent of the phase separation observed between noisy (i.e. active) and passive particles in colloidal mixtures (43, 44). Thus, the surface fluctuations could direct sorting independently of cortical tension.

Our data suggest that ERM-based regulation of effective membrane tension could be an important and previously overlooked player in tissue organisation. Indeed, changing ERM levels directly affected sorting both in vivo (Fig 5d-e) and in cell aggregates (Fig 5g-j). However, our analysis also strongly suggests that it is not necessarily absolute levels of membrane tension that is important for sorting, but its dynamics and spatial heterogeneity, which correlate with surface fluctuations and thus regulate spatial segregation of cell lineages. Future work will be needed to disentangle what other molecular players control the levels of heterogeneities in membrane tension, and the organisation of the underlying actin cortex, in the developing embryo.

Ultimately, our discovery that differences in noise at the cell surface, or surface fluctuations, drive lineage segregation in the mouse blastocyst provides new insight into tissue self-organisation in the early embryo. It will be interesting to investigate how cell surface dynamics influence other processes of self-organisation across organisms, including tissue morphogenesis and tumorigenesis.

## ACKNOWLEDGEMENTS

We thank H. Niwa for Dox regulatable PB vector, G. Charras for EzrinT567D cDNA, K Jones for tdTomato ESCs and general lab managements, M. Kinoshita for pPB-CAG-H2B-BFP plasmid, P. Humphreys for assistance with imaging, G. Chu, P. Attlesey and staff for animal husbandry, C. Mulas for critical feedback on the manuscript, T. Boroviak for single-cell RNA-seq, the EMBL Genomics Core Facility for sequencing. We also thank all Stem Cell Institute staff for helping us to maintain our productivity. This work was financially supported by the Wellcome Trust, Medical Research Council, Leverhulme Trust (RPG-2014-080), BBSRC (BB/Moo4023/1), a Next Generation fellowship awarded to S.A. by the Centre for Trophoblast Research, British Council and JSPS Overseas Research Fellowships awarded to A.Y.. MRC Career Development Award (G1100312/1) to K.F.

## AUTHOR CONTRIBUTIONS

AY and ECS performed embryology and wet laboratory experiments; DC, AY, and SA performed AFM experiments; GS was responsible for bioinformatics and data analysis; CR, RB, and KC were responsible for computational simulations; ECS, AY and HDB performed membrane tension measurements; AY and IMA performed migration assay; ECS, AY and RP performed dSTORM imaging, KC and AY were responsible for quantitative analysis of imaging data; AY and KC wrote the paper. KC, KF, JN and EP discussed the data and assisted with manuscript preparation. KC and JN initiated and supported the research.

## COMPETING FINANCIAL INTERESTS

The authors declare no competing financial interests.

## Notes

### Competing Interest Statement

The authors have declared no competing interest.

