## Supplementary materials for "Cell surface fluctuations regulate early embryonic lineage sorting"

#### Materials and Methods

##### Mouse strains and embryo collection

Mice used were intercrosses of *Pdgfra*$^{H2B\text{-}GFP/+}$ (*1*), in which a cassette containing human H2B fused to enhanced green protein (H2B-GFP) was targeted to the *Pdgfra* locus, and first filial generation (F1) hybrids (C57BL/6JxCBA/J) (Charles River), homozygous mTmG [Gt(ROSA)26Sor$^{tm4(ACTB-tdTomato,\ -EGFP)Luo}$, mTmG$^{+/+}$] (*2*) or CD-1 (Charles River). All embryos used in this study were obtained from natural mating. Embryo staging was based on the assumption that, on average, mating occurred at midnight so that at midday, the embryos were assigned E0.5. Embryos were flushed at the relevant stages from oviducts (eight-cell stage embryos) or uterine horns (blastocysts) using flushing and holding media (M2, Sigma). PrE cells can be visualised with *Pdgfra*$^{H2B\text{-}GFP/+}$ reporter which we used as additional criteria to classify embryo stages (*3*, *4*). EPI and PrE have not segregated at E3.5 and E3.75 stages. EPI and PrE have segregated at E4.0 and E4.5. GFP positive PrE were clearly seen to form one layer faced with a blastocoel. The sex of embryos and the ages of mice using mating were not concerned in this study. This research has been regulated under the Animals (Scientific Procedures) Act 1986 Amendment Regulations 2012 following ethical review by the University of Cambridge Animal Welfare and Ethical Review Body (AWERB). Use of animals in this project was approved by the ethical review committee for the University of Cambridge, and relevant Home Office licences (Project licence No. 80/2597 and No. P76777883) are in place.

##### Isolation of ICMs from embryos, and single-cell dissociation of ICMs

Embryo and cell manipulations were carried out under a dissecting microscope (Leica Microsystems). The zona pellucida was removed using acid Tyrode's solution (Sigma). Blastocysts from E3.5-E4.5 were subjected to immunosurgery as previously described (*5*). In brief, blastocysts were incubated for 45-60 minutes in a 1:5 dilution of anti-mouse rabbit serum (Sigma) in N2B27, washed in N2B27 and further incubated for 30-60 minutes in a 1:5 dilution of rat serum (in-house) in N2B27 for the complement reaction. The ICM was subsequently cleaned from residual trophectoderm with a narrowly fitting glass pipette. Single-cell dissociation of ICMs was performed in a 1:1 mixture of Accutase (PAA) and 0.025% trypsin (Invitrogen) plus 1% chick serum (Sigma). Cells were dissociated by repetitive using blunted microcapillaries (Global Scientific or Harvard apparatus) and washed in Blast (Origio) or N2B27. N2B27 media were prepared as described (*6*). Briefly, 1:1 Dulbecco's Modified Eagle's Medium/Nutrient Mixture F-12 Ham (DMEM/F-12; Sigma) and Neurobasal media (Gibco), N2 (in-house) and B27 (Thermo Fisher Scientific) additives, 2 mM L-glutamine (Thermo Fisher Scientific), and 100 mM 2-mercaptoethanol (Sigma) were supplemented.

**cDNA amplification and synthesis from single cells**

Three E3.75 *Pdgfra*$^{H2B\text{-}GFP/+}$ positive embryos obtained from intercrossing of *Pdgfra*$^{H2B\text{-}GFP/+}$ and F1 hybrids were used. We collected embryos, which had salt-and-pepper GFP positive cells distribution in ICM and proper GFP intensity. ICM cells were dissociated from isolated ICM by immunosurgery. Dissociated single cells were transferred immediately into Smart-Seq2 single-cell lysis buffer and immediately frozen on dry ice. Smart-Seq2 library was prepared as originally describe (*7*) and sequenced on the Illumina HiSeq2000 platform (150 base, paired end).

**RNA-seq data processing**

Sequencing data of single-cell mouse embryo profiling study (accession SRP110669 (*8*)) was downloaded from the European Nucleotide Archive (*9*). Mus musculus GRCm38.87 gene annotation was used together with mm10 genome version. Alignments to gene loci were quantified with htseq-count (*10*) based on annotation from Ensembl 87. Sequencing libraries with fewer than 500K mapped reads were excluded from subsequent analyses. Read distribution bias across gene bodies was computed as the ratio between the total read spanning the 50th to the 100th percentile of gene length, and those between the first and 49th. Samples with ratio >1.5 were not considered further. Stage-specific outliers were screened by principal component analysis.

**Transcriptome analysis**

Principal component and cluster analyses were performed based on $\log_2$ fragments per kilobase of exon per million mapped fragments ($\log_2$FPKM) values computed with the Bioconductor packages *DESeq2* (*11*), *Sincell* (*12*) or *FactoMineR* (*13*) in addition to custom scripts. Differential expression analysis was performed with scde (*14*), which fits individual error models for the assessment of differential expression between sample groups. Pseudotimes were computed using R monocle package (*15*). For global analyses, genes that registered zero counts in all single-cell samples in a given comparison were omitted. Euclidean distance and average agglomeration methods were used for cluster analyses. Expression data are available upon request. Ensembl 87 annotation was used to download specific actin cytoskeletal genes using biological process name as a keyword.

**Selection of high-variability genes**

Gene exhibiting the greatest expression variability (and thus contributing substantial discriminatory power) were identified by fitting a non-linear regression curve between average $\log_2$ FPKM and the square of the coefficient of variation. Thresholds were applied along the $x$-axis (average $\log_2$ FPKM) and $y$-axis (log squared coefficient of variation [$CV^2$]) to identify the most variable genes. As actin-cytoskeleton related genes, we selected 6899 genes (Supplementary Data Table1). Amongst them, 152 genes were highly modulated in ICM cells through E3.5 to E4.5 blastocysts and use these genes for actin-cytoskeleton related genes principal component and cluster analyses (Fig. 1e, Supplementary Data Table2).

**Embryo and ICM culture**

Embryos and isolated ICMs were cultured in Blast, KSOM (Millipore) or N2B27 in an organ culture dish (Falcon) culture or in single-drop cultures under mineral oil (Sigma) in a humidified incubator at 37°C with 5% $CO_2$. Embryo culture media were buffered in the incubation chamber for at least 30 minutes before embryos and ICM culture.

**Live imaging of isolated ICMs**

Isolated ICMs were transferred to an embryo immobilization chip (Dolomite Centre Ltd). The spinning disk microscope (Andor Revolution XD System [ANDOR] with a Nikon Eclipse Ti microscope [Nikon]) was used for taking images. 17 z-stacks per time step every 30 minutes were taken, with three channels (488 nm excitation for $Pdgfra^{H2B\text{-}GFP/+}$ reporter, 561 nm excitation for membrane [mTmG] and bright field). An Andor 85 camera recorded images with magnification through a CFI Plan Fluor ×40/1.3 oil objective (Nikon) with Cargille microscope immersion oil (Cargille Labs). Each experiment was set up using Andor IQ Software. Each image collected data in 502 × 501 (width × height) pixels. The microscope is equipped with an incubation chamber to keep the sample at 37°C and 7% $CO_2$. Images were processed using Fiji (*16*).

**Doublet formation**

**Forming doublets:** E3.75 $Pdgfra^{H2B\text{-}GFP/+}$ positive embryos obtained from intercrossing of $Pdgfra^{H2B\text{-}GFP/+}$ and F1 hybrids were used. Two isolated single ICM cells were put together by gently blowing the surrounding medium through a microcapillary in the micro drop of Blast under the mineral oil and incubated for 30 minutes at 37°C and 5% $CO_2$. When two isolated ICM cells come into contact, the contact grows until equilibrium is attained. Doublets were transferred in Blast drop in the presence of 0.01% CellMask™ Deep Red Plasma membrane Stain (Thermo Fisher Scientific) to visualise membrane under the mineral oil in a glass-bottom dish (MatTek) coated with poly-D-lysine (PDL, Millipore) (see Extended Supplementary information). For the coating, the dishes were coated with small drops of 50 µg/ml PDL for at least one hour at room temperature. The drops were washed three times with the media used for imaging and covered with mineral oil. Before transferring doublets to Blast drop, the drops under mineral oil were buffered in the incubation chamber for at least 30 minutes. Confocal images were acquired using a Leica TCS SP5 (Leica Microsystems) confocal microscope. Optical section thickness was 0.99 µm. A HC PL APO 40×/1.30 Oil CS2 (Leica) with immersion oil (Leica) was used. Whole doublet images from bottom to top were taken with three channels (488 nm excitation for *Pdgfra*$^{\mathrm{H2B\text{-}GFP}}$ reporter, 647 nm excitation for membrane [CellMask™ Deep Red Plasma membrane Stain] and bright field). The microscope is equipped with an incubation chamber to keep the sample at 37°C and 7% $CO_2$.

**External contact angle measurement:** The external contact angles at the middle section of the doublets were measured by using the angle tool of Fiji. The average of both sides of the external contact angles was used as the external contact angle $\theta_e$. The mean GFP intensity of *Pdgfra*$^{H2B\text{-}GFP}$ was quantified using Fiji and used to classify the lineage of each ICM cell. The top and bottom 40% of cells with strong GFP intensities (GFP$^{\mathrm{high}}$ cells and GFP$^{\mathrm{low}}$ cells) were considered as pPrE cells and pEPI cells, respectively, and used for this analysis. The doublets that were not horizontal to the dish, mitotic cells, dead cells and blebbing cells at the interface were excluded from the analysis.

**CS3D is described in full in the Extended Supplementary Information and** (*17*)**.**

**Cell preparation and experimental setup for ICM cell/aggregate surface fluctuation analysis:** mTmG$^{+/-}$*Pdgfra*$^{H2B\text{-}GFP/+}$ positive embryos obtained from intercrossing of *Pdgfra*$^{H2B\text{-}GFP/+}$ and mTmG$^{+/+}$ were used. For single ICM cell membrane dynamics study, isolated single ICM cells were transferred in the Blast drops under mineral oil on a PDL-coated glass-bottomed dish. The cells were kept for 15 minutes in a humidified incubator at 37°C and 5% $CO_2$ and subsequently kept for 15 minutes in an imaging chamber at 37°C and 7% $CO_2$ before imaging for purposes of equilibration. For the cytokine and inhibitor experiments, 25 ng/ml FGF2 (in-house), 1 μM PD03 (abcr) or 0.01% DMSO (Thermo Fisher Scientific) was added to Blast drops. For ICM aggregates membrane dynamics study, small depressions were indented on a 60-mm dish (Thermo Fisher Scientific) lid by a sterilised aggregation needle (BLS Ltd) and covered with Blast drops (one depression per one drop) overlaying with mineral oil. Three isolated E3.75 ICMs were disposed to make a triangle in a small depression and kept for one hour in a humidified incubator at 37°C and 5% $CO_2$. Aggregated ICMs were transferred in the Blast drops under mineral oil on a PDL-coated glass-bottomed dish. Before transferring ICM cells or aggregated ICMs to Blast drops, the drops under mineral oil were buffered in a humid incubator at 37°C and 5% $CO_2$ for at least 30 minutes. Live images were acquired using Leica TCS Sp5 Confocal microscopy on the single middle z-slice through the cell every ten seconds for ten minutes with three channels (488 nm excitation for *Pdgfra*$^{H2B\text{-}GFP/+}$ reporter, 567 nm excitation for membrane [mTmG] and bright field). For ICM aggregates membrane dynamics study, live images were taken on the several z-slice through the aggregates every 20 seconds for ten minutes.

**Cell lineages classification:** The mean intensity of *Pdgfra*$^{\mathrm{H2B\text{-}GFP}}$ was measured by Fiji. ICM cell lineages were determined by their GFP signal. The E3.5 cells with GFP positive were classified as pPrE. The E3.5 cells with GFP negative were classified as ICM/pPrE cells. The E3.75 cells with the top 40% of GFP intensity were classified as pPrE, the E3.75 cells with the bottom 40% of GFP intensity were classified as pEPI. The E4.0 cells with GFP positive were classified as PrE. The E4.0 cells with GFP negative were classified as EPI. Mitotic cells judging by H2B-GFP morphologies were removed from the analysis.

**Quantification of surface fluctuations:** Each cell's live imaging data was cropped and registered with StackReg plugin (*18*) using Fiji. The centroid of the first images was used as the centre of the new coordinates system and linear interpolation (See Fig. 3b and Extended Supplementary Information). The position of the cell membrane (tdTomato signal of mTmG) was identified and converted from Cartesian coordinates into polar coordinates. The boundary coordinate plots of the radius versus the angular coordinate ($\theta$) were detrended to set the average radial value is zero. This normalises for differences in the cell size and controls for small fluctuations of the focal plane in the *z*-axis. The variation in time ($V_{\mathrm{T}}$) and) was calculated using equation (1), $\rho$: distance from the centre of a cell.

$$V_{\mathrm{T}} = \overline{\left(SD\left(\{\rho_{\mathrm{j}}(t_1), \rho_{\mathrm{j}}(t_2), \dots, \rho_{\mathrm{j}}(t_{\mathrm{N}})\}\right)\right)_{\mathrm{j}=1:\mathrm{M}}} \quad (1)$$

Additional information about this analysis is included in Supplemental Information.

**ICM aggregate surface fluctuation analysis**

Surface fluctuations on outside cells were scored from 1 to 5, with 1 being no observable fluctuations and 5 being significant observable fluctuations, using only mTmG time-lapse images so as not to identify each cell lineages. A high score indicates that the cell has a dynamic cell membrane movement. This analysis was done single-blind by AY and KC.

**ES cells culture**

ES cells were routinely maintained on 0.1% gelatine (Sigma)-coated 6-well plates (Falcon) in 2i+LIF media(*6*), which contains N2B27 medium supplemented with 1 μM PD03 and 3 μM CHIR99021 with 10 ng/ml LIF (in-house). Cells were passaged every three days, using Accutase disassociation.

**Generation of H2B-BFP and tdTomato ES cells**

tdTomato ES cells were generated from the embryo crossed (Rosa)26Sor$^{tm9(CAG\text{-}tdTomato)Hze}$ mice (JAX#007909 (*19*)) with R26Cre$^{ER}$ mice (JAX#004847 (*20*)). 500 nM 4-Hydroxytamoxifen (Sigma) was added to the ES cells and tdTomato-positive ES cells were expanded. For H2B-BFP ES cells generation, 0.8 μg of pPB-CAG-H2B-BFP-IRES-Neo (kindly gifted by M Kinoshita) and 0.4 μg of pPy-CGA-PBase were transfected into E14Tg2A (E14) ES cells using Lipofectamine 2000 (Thermo Fisher Scientific). Following the drug selection with 400 μg/ml G418 (Thermo Fisher Scientific), BFP-positive colonies were picked and expanded.

**RNA interference**

$1x10^4$ cells tdTomato ES cells were seeded on 0.1% gelatine coated 24-well plates. Cells were transfected with 15 μM of either the targeting (5 μM each SMARTpool siGENOME Mouse Ezrin[M-046568-01-0005], Moesin[M-044428-01-0005], Radixin[M-047230-01-0005]; Dharmacon) or control siRNA (D-001210-02-05, Dharmacon) with Lipofectamine RNAiMAX Transfection Reagent (Thermo Fisher Scientific) for overnight. The media were changed the next day. The transfected cells were utilised for cell shape imaging or chimaera assay. Knockdown efficiency was checked using RT-qPCR (see Extended Supplementary Information).

**Generation of chimaeras**

ES cells (three-five cells per embryo) were injected into early blastocysts (E3.5) via a laser-generated perforation in the zona pellucida using XYClone (Hamilton Thorne Biosciences). Injected embryos were cultured in Blast for a half-day, the equivalent of E4.0 blastocysts at 37°C and 5% $CO_2$.

**Immunofluorescence staining**

Embryos were fixed with 4% paraformaldehyde (PFA; Thermo Fisher Scientific) in Phosphate buffered saline (PBS; Sigma) at room temperature for 15 minutes. Then, the samples were rinsed in PBS containing 3 mg/ml polyvinylpyrrolidone (PBS/PVP; Sigma), permeabilised with PBS/PVP containing 0.25% Triton X-100 (Thermo Fisher Scientific) for 30 minutes. Blocking was performed with an embryo blocking buffer comprising PBS containing 0.1% bovine serum albumin (BSA; Sigma), 0.01% Tween20 (Sigma) and 2% donkey serum (Sigma) at 4°C for 2-3 hours. Samples were incubated in the blocking buffer with 500 ng/ml 4',6-diamidino-2-phenylindole (DAPI; Invitrogen) at room temperature for one hour in the dark. They were rinsed three times in an embryo blocking buffer for 15 minutes ~ each, then transferred in small drops of blocking buffer on a PDL-coated glass-bottom dish under the mineral oil and taken images. Whole staining process was performed on Pyrex 9 depression spot plate (Corning).

For pERM staining, pEPI and pPrE cells seeded on a PDL-coated 10-well slide glass (TF1006, MTSUNAMI) were fixed with 4% PFA in cytoskeletal stabilizing buffer (CSB; 10 mM MES pH6.1, 138 mM KCl, 3 mM $MgCl_2$, 2 mM EGTA) containing 4.5% w/v sucrose (Sigma) and 0.2% Triton X-100 at 37°C for six minutes. Subsequently, the samples were fixed with 4% PFA in CSB containing 4.5% w/v sucrose at 37°C for 14 minutes. The fixation buffers were pre-warmed at 37°C before use. Then, the samples were rinsed in PBS for twice, permeabilised with PBS containing 0.1% Triton-X at room temperature for ten minutes. Blocking was performed with a buffer comprising PBS containing 2% FBS, 2% BSA and 0.1% Triton-X at room temperature for 45 minutes. Primary antibodies were diluted in blocking buffer, and samples were rinsed with the appropriate antibody solution once and incubated with the antibody solution at 4°C overnight. They were rinsed five times using PBS containing 0.1% Triton-X for five minutes ~ each. Secondary antibodies were diluted in blocking buffer, and samples were incubated in the appropriate antibody solution at room temperature for one hour in the dark. They were rinsed five times in PBS containing 0.1% Triton-X for five minutes ~ each. 10 µl of Vectashield (Vector Laboratories) was added to each well, and coverslips were mounted and sealed with nail varnish. Primary and secondary antibodies were listed in Supplementary Data Table 3.

**Imaging**

For embryos, isolated ICMs, doublets of ICM cells, ES cells and ES cell aggregates imaging, samples were transferred to the drops on PDL-coated glass-bottom dishes and taken images using a Leica TCS SP5 or ZEISS LSM880 (Carl Zeiss Microscopy) confocal microscope. For live imaging of ICM aggregates, doublets of ICM cells and ES cells, SP5 was used with an incubation chamber to keep the sample at 37°C and 7% $CO_2$. For the quantification of fluorescence in single cells, a set of experiment images were acquired using the same microscope with the same setting on the same day. Images were processed using Fiji.

**Surface fluctuation analysis of ES cells**

For CA-EZR-IRES-mCherry ES cells were plated for 24 hours in N2B27+2i+LIF media at $2 \times 10^4$ cells/cm$^2$, and the media were changed to N2B27+2i+LIF media with or without 1 µg Dox. Then, cells were dissociated into a single-cell suspension using Accutase and suspended in 100 µl each culture media with 0.01% CellMask™ Green Plasma membrane Stain (Thermo Fisher Scientific). The media were buffered in the incubation chamber for at least 30 minutes before suspension. The analysis was performed as outlined in Supplementary information Fig.4.

**Generation of Dox-inducible EzrinT567D (CA-EZR) ES cells**

Human EzrinT567D (constitutive active form of Ezrin; CA-EZR) cDNAs were a kind gift from G. Charass. They were inserted into pPB-CMV-HA-pA-IRES-Neo (kindly gifted by H. Niwa) using In-Fusion (Clontech) to generate pPB-Tet-CA-EZR. 0.8 µg of pPB-Tet- CA-EZR was transfected with 0.8 µg of pPB-CAG-rtTA-IRES-Neo (kindly gifted by A Smith [Addgene plasmid #606612] (*21*))and 0.4 µg of pPy-CGA-PBase using Lipofectamine 2000 into E14 ES cells. The cells were harvested from a 0.1% gelatin-coated 6-well plate in 2i+LIF media. After 48 hours, 400 µg/ml G418 was added to ES cells and colonies were selected. After one week of G418 selection, clones were manually picked, dissociated, then split into a 96-well plate (Corning). ES cells were cultured with 2i+LIF in the presence or absence of 1 µg/ml Dox (Sigma). Clones that with no mCherry signal in the absence of Dox and high mCherry signal in the presence of Dox were chosen by eyes.

**CA-EZR overexpression and imaging**

For CA-EZR overexpressing experiments, cells were plated for 24 hours in 2i+LIF media at $2 \times 10^4$ cells/cm$^2$, and the media were changed to 2i+LIF in the presence of 1 μg Dox (Sigma). For the imaging of pERM in single CA-EZR cells, cells were cultured in 2i+LIF in the presence of 1 μg Dox for 24 hours prior to being plated onto imaging dishes (μ-Dish 35 mm ibidi dish [81156]). After six hours, the cells were fixed using 4% in CBS for 15 minutes and then permeabilised in with 0.1% Triton in CBS. Blocking was then performed using 2% FBS, 2% BSA in PBS with 0.1% Triton-X for 45 minutes. Cells were then incubated for 90 minutes with the primary antibody in the same buffer as used for blocking. Cells were then washed three times for five minutes with PBS containing 0.1% Triton-X. Secondary antibodies were added together with Alexa Fluor™ 647 Phalloidin for one hour in the same buffer as used for blocking and primary antibody incubation. Cells were washed with PBS containing 0.1% Triton-X three times for five minutes before a final wash in PBS.

**ES cells aggregation**

ES cells harvested on tissue culture plates were detached using Accutase. The cells were suspended in FBS (GE Healthcare Life Sciences) containing suspension media, and cell concentration was determined. Suspended cells were centrifuged at 1400 rpm for three minutes, pelleted and re-suspended in 2i+LIF media in the absence or presence of 1 μg/ml Dox. H2B-BFP and CA-EZR-IRES-mCherry ES cell lines were mixed well in a universal tube (Scientific Laboratory Supplies Ltd) at 1:1 ratio and then plated 200 μl per well in round-bottomed low-adhesion 96-well plates (CELLSTAR) with 300 cells per well. After 30 hours culture, aggregated ES cells were collected by mouth pipette, briefly rinsed in PBS, fixed with 4% PFA for 20 minutes at room temperature, rinsed in PBS/PVP, and then incubated briefly in increasing concentrations of Vectashield before mounting on glass slides (Thermo Fisher Scientific) in small drops of concentrated Vectashield. Subsequently, coverslips with Vaseline (Unilever) spacer were mounted and sealed with nail varnish. FBS containing suspension media was composed of Glasgow's minimum essential medium (GMEM) with 10% batch-tested FBS, 1× MEM non-essential amino acids (NEAA; Sigma), 1 mM sodium pyruvate (Sigma), and 1 mM L-glutamine, 0.1 mM 2-mercaptoethanol. For calculation of the average radial distance from the centre, *R,* we assessed the distribution of mCherry levels in the cells. The bottom half of the distribution was assumed to be the control cells and was discarded. In the top 50% expressing cells, the distributions were split into a bottom 1/3, a middle 1/3, and a top 1/3. These were considered, respectively, as the low-, mid-, and high-expressing cells. These were then binarized and used in the formula shown in Fig. 4h to find *R*.

**Data availability**

Codes and any other data that support the findings of the study are available from the corresponding authors upon reasonable request.

**Statistical analysis**

For all statistical analysis in the paper, unless otherwise indicated in corresponding figure legends, n-way ANOVA was used to calculate P-values to establish significant changes between any two means. Details of each ANOVA stated in figure legends: for example, variables used for the n-way ANOVA could be 'replicate number', 'cell type' (e.g. EPI and PrE) and when relevant 'small molecule treatment'. There was usually a significant interaction effect for 'experiment number' due to expected experimental variability in embryo work, so this $p$-value is suppressed throughout the paper and the $p$-value reported is for 'cell type' or 'small molecule treatment' depending on the experiment. The midline is mean of overall experiments, and error bars represent standard deviation over all experiments. For Fig. 2d, standard deviations were calculated as the square root of the sum of the squares of the standard deviation of each experiment divided by the number of experiments.

**Materials and Methods References**

1. T. G. Hamilton, R. A. Klinghoffer, P. D. Corrin, P. Soriano, Evolutionary divergence of platelet-derived growth factor alpha receptor signaling mechanisms. *Mol. Cell Biol.* **23**, 4013–4025 (2003).

2. M. D. Muzumdar, B. Tasic, K. Miyamichi, L. Li, L. Luo, A global double-fluorescent Cre reporter mouse. *Genesis*. **45**, 593–605 (2007).

3. B. Plusa, A. Piliszek, S. Frankenberg, J. Artus, A.-K. Hadjantonakis, Distinct sequential cell behaviours direct primitive endoderm formation in the mouse blastocyst. *Development*. **135**, 3081–3091 (2008).

4. J. B. Grabarek *et al.*, Differential plasticity of epiblast and primitive endoderm

precursors within the ICM of the early mouse embryo. *Development*. **139**, 129–139 (2012).

### Supplementary Figures

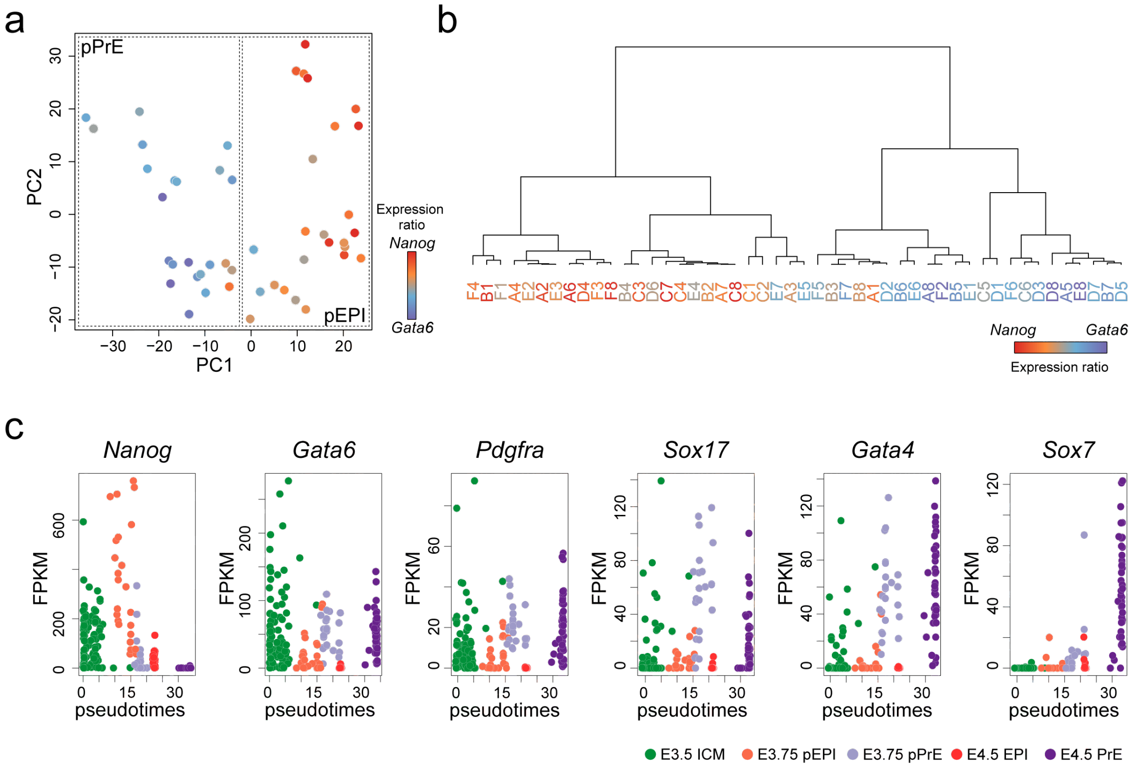


**Fig. S1: Profiling of E3.75 pEPI and pPrE cells.**
**(a)** PCA of E3.75 single cells computed with highly variable genes (n = 3259, $\log_2$ FPKM> 0.5, $\log CV^2$> 0.25) **(b)** Dendrogram of E3.75 ICM cells based on variable genes for E3.75 stage; expression coloured according to the ratio of *Nanog* to *Gata6* expression. **(c)** Expression profiles of selected genes ordered by pseudo time scale.

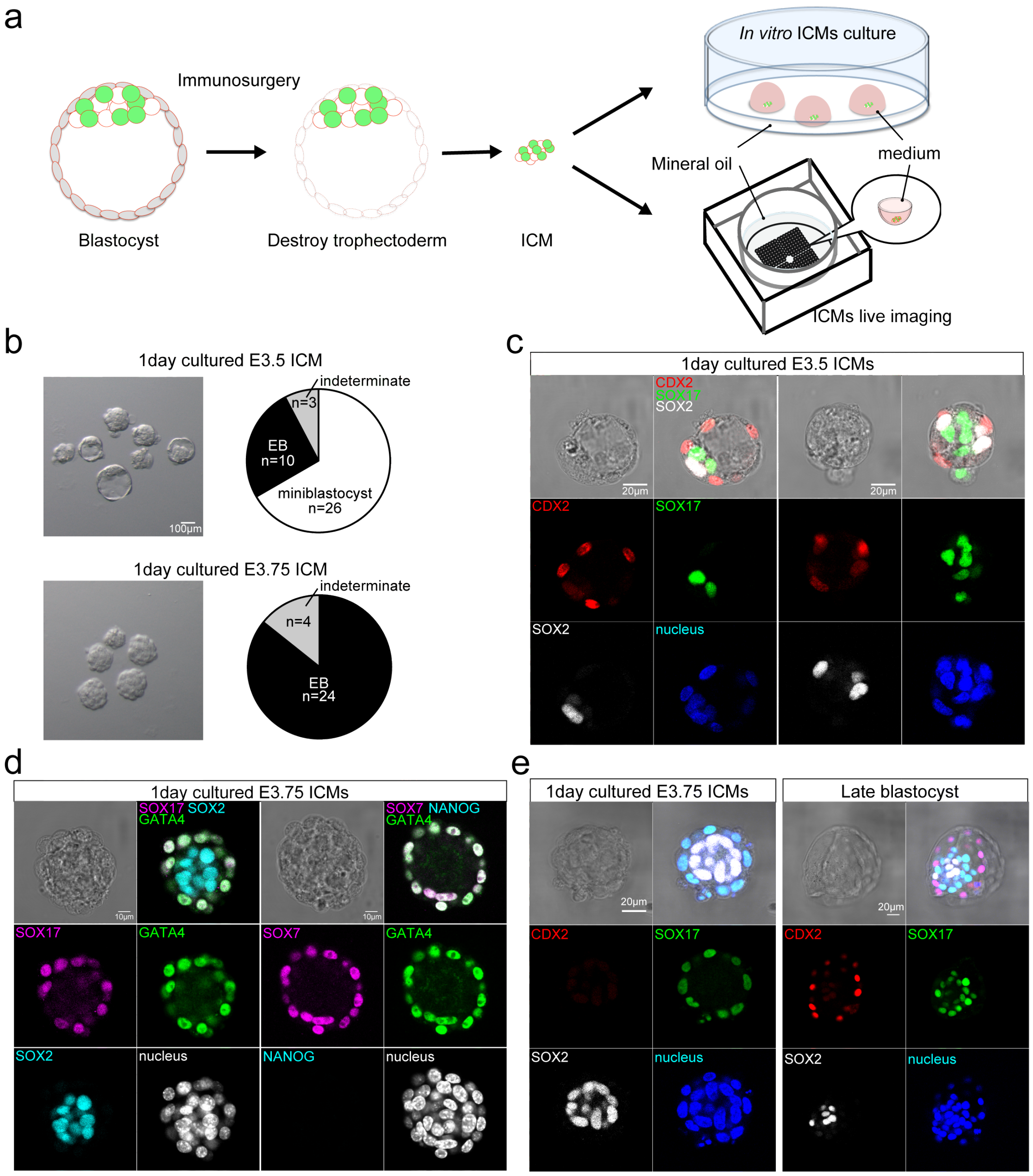


**Fig. S2: pEPI and pPrE begin to segregate at approximately E3.75.**
**(a)** Schematic images of ICM isolation from blastocyst and culture. Sequential images of isolated E3.75 mTmG$^{+/-}$*Pdgfra*$^{H2B\text{-}GFP/+}$ ICM culture was performed in an embryo immobilisation chip using spinning disk confocal microscopy. **(b)** Bright-field images and the proportion of miniblastocysts, i.e. ICMs with a cavity embryoid bodies (EBs) and indeterminate morphologies of isolated E3.5 and E3.75 ICMs cultured for one day. Note that E3.75 cultured ICMs lose the capacity to form a mini-blastocyst. **(c)** Representative images of one day cultured isolated E3.5 ICMs. EPI marker, SOX 2, PrE maker, SOX17 and TE marker, CDX2 were expressed in one day cultured E3.5 ICMs. **(d)** Immunofluorescence staining of isolated E3.75 ICMs cultured for one day from *Pdgfra*$^{H2B\text{-}GFP/+}$. SOX2 is an EPI marker, and NANOG is an early EPI marker. SOX17 and GATA4 are PrE markers. SOX7 is late PrE marker. **(e)** Representative images of one day cultured isolated E3.75 ICMs and late-stage blastocyst. EPI marker, SOX 2 and PrE maker, SOX17 were expressed in one day cultured E3.75 ICMs. TE marker, CDX2 was expressed in late-stage blastocyst but not in one day cultured E3.75 ICMs, indicating that, unlike E3.5 cultured ICMs, E3.75 cultured ICMs lose the capacity to make TE.

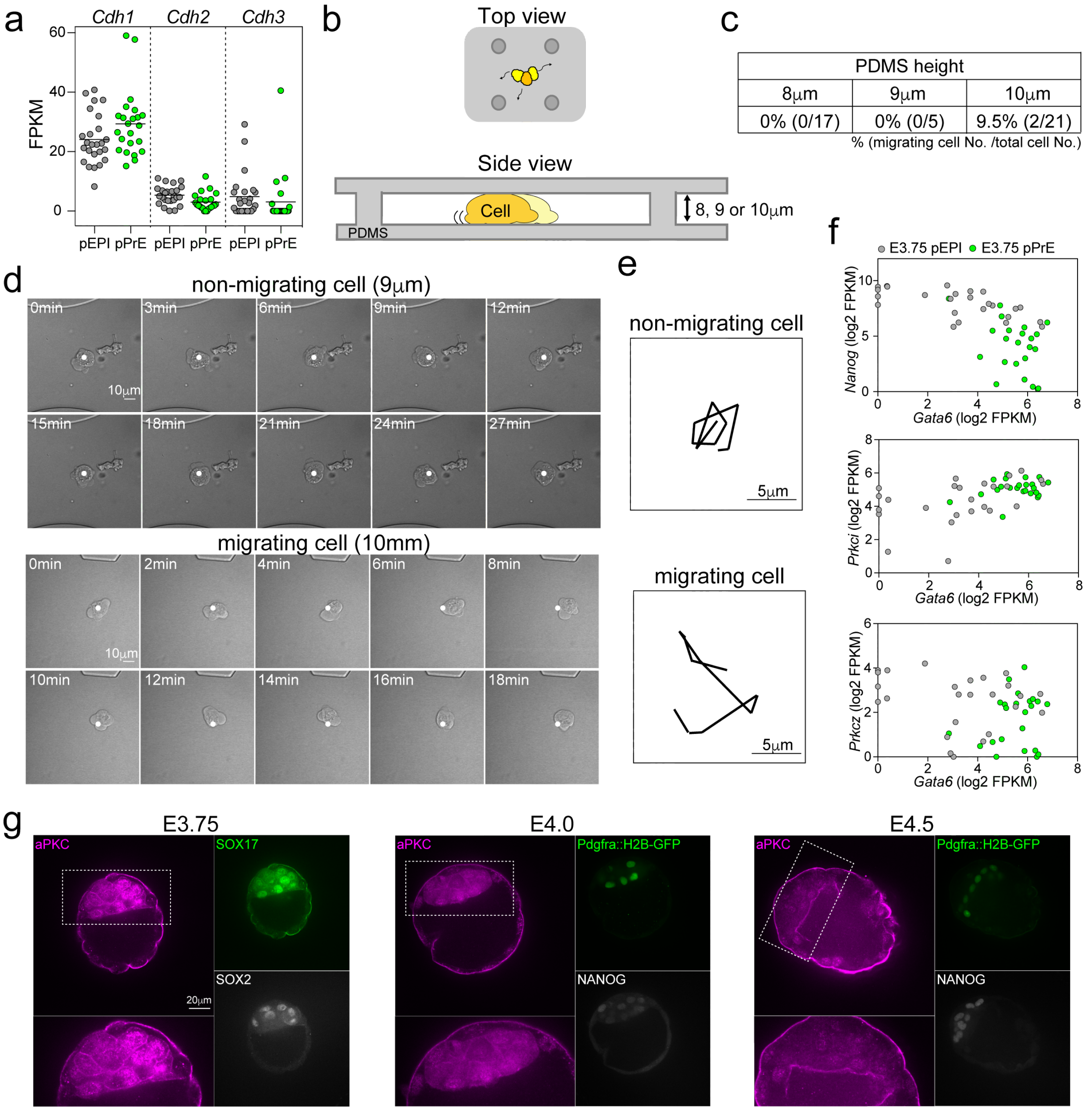


**Fig. S3: Differences in cell-cell adhesion, polarity or migration do not adequately explain cell sorting.**
**(a)** The mRNA expression level of *cdh1* (E-cadherin), *cdh2* (N-cadherin) and *cdh3* (P-cadherin) in E.3.75 pEPI and pPrE, indicating very little differential expression of adhesion factors between pPrE and pEPI. **(b)** Schematic of a polydimethylsiloxane (PDMS) confinement device for testing cell migration potential in confinement. **(c)** The proportion and number of E3.75 ICM cells migrating under 8, 9 or 10 μm height confinements, indicating that these cells have very little capacity to undergo confined migration. **(d)** Representative images of time series of non-migrating and migration E3.75 ICM cells using 9 μm or 10 μm height confinement. **(e)** Trajectories of the two E3.75 ICM cells in (d) during ten sequential frames. Note that 12 μm channels were also attempted but did not confine the cells. **(f)** Correlation plot of the mRNA expression levels of *Gata6* (PrE marker) and either *Nanog* (EPI marker), or the polarity factors *Prkci* (aPKCλ) and *Prkcz*(aPKCζ), in E3.75 pEPI and pPrE, indicating that differential expression of polarity factors has not yet been firmly established at E3.75. **(g)** Representative images of aPKC, SOX2 and SOX17 or Pdgfra::H2B-GFP in E3.75, E4.0 and E4.5 blastocysts. Note that at E3.75 and E4.0 aPKC appears to be primarily cytoplasmic and not yet localised to the membrane, as it is at E4.5.

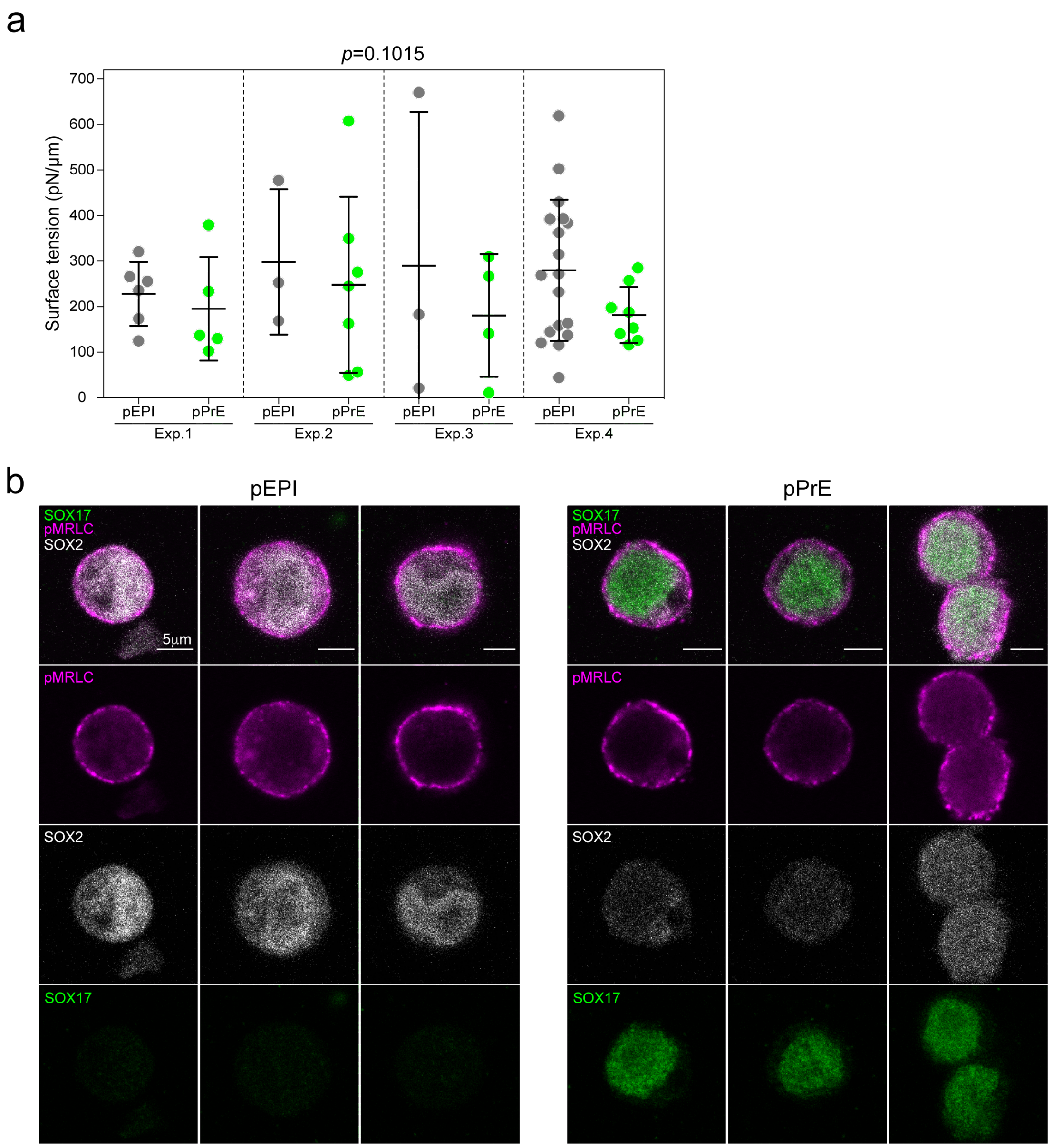


**Fig. S4: No reproducible distinction between pEPI and pPrE in surface tension.**
**(a)** The surface tension of dissociated pEPI and pPrE from E3.75 *Pdgfra*$^{H2B\text{-}GFP/+}$ embryos measured using an atomic force microscope (AFM), using the technique presented in *Chugh P et. al. Nat Cell Biol 2017*. P-value was calculated by 2-way ANOVA using cell type and experimental replicate as variables. **(b)** Representative images of pMRLC (magenta), a proxy for cytoskeletal tension, in E3.75 pEPI and pPrE, further indicating that there is little difference in surface tension between pEPI and pPrE. SOX2 is an EPI marker. SOX17 is a PrE marker.

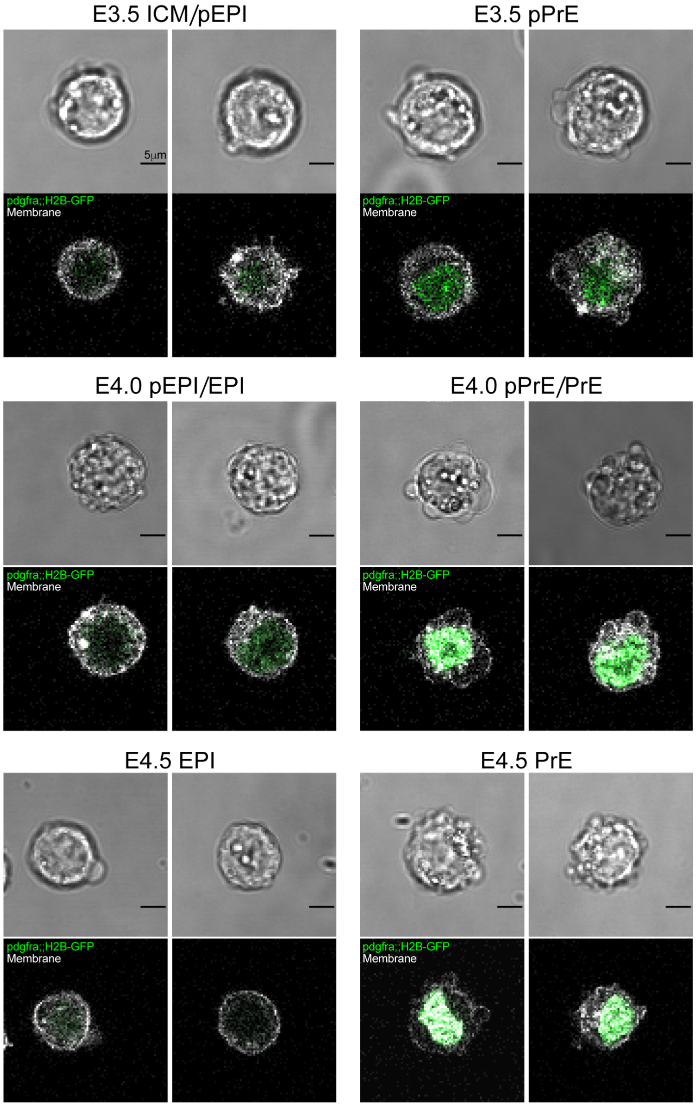


**Fig. S5: In the ICM, the PrE lineage blebs more than the pEPI lineage.**
Representative images of isolated ICM cells from *Pdgfra*$^{H2B\text{-}GFP/+}$ E3.5, E4.0 and E4.5 blastocysts, indicating that at all stages the PrE lineage and its progenitors exhibit more blebbing than the EPI lineage and its progenitors. Note, however, that at E4.5, the blebs tend to present as smaller blebs, ruffles or protrusions. Pdgfra starts to be expressed in pPrE at the early blastocyst stage. Pdgfra negative E3.5 ICM cells are either pEPI or ICM cells, which have not specified their lineages yet. E4.0 ICM cells may contain pEPI, pPrE, EPI and PrE.

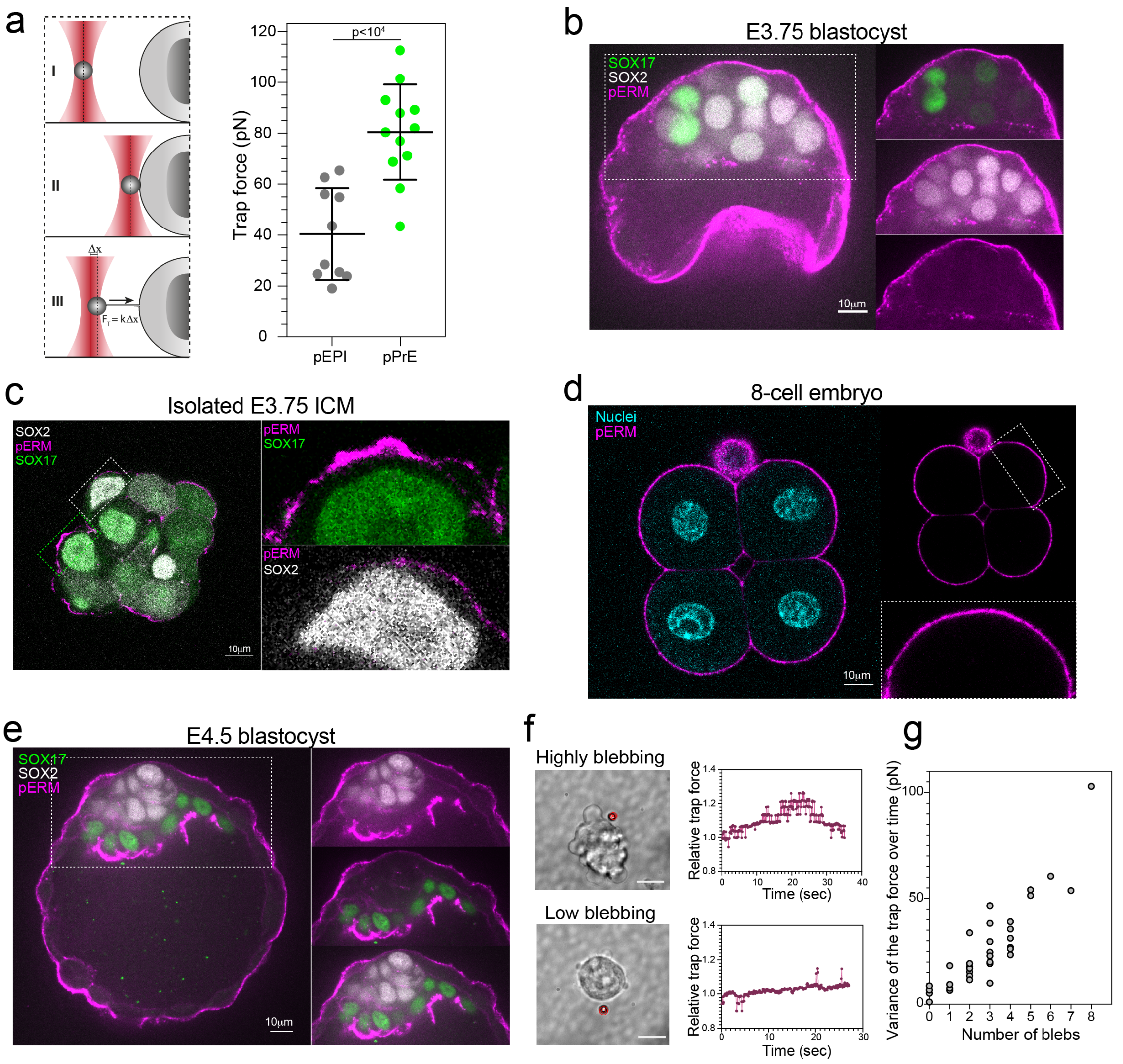


**Fig.S6: pERM and membrane tension variability is likely responsible for enhanced surface fluctuations in PrE lineage.**
**(a)** Schematic of optical tweezers to measure membrane tension in a cell. The membrane tension, as measured by trap force, of E3.75 pEPI and pPrE isolated from *Pdgfra*$^{H2B\text{-}GFP/+}$ embryos. P-value was calculated by 1-way ANOVA. **(b-e)** Representative images of pERM in E3.75 blastocyst, E3.75 isolated ICM, 8-cell embryos and E4.5 blastocyst, indicating that pERM is much more variable on the surfaces of ICM cells at E3.75 than other stages and lineages in the early embryo. **(f)** Left, representative images of membrane tension measurement of an ES cell that is highly blebbing (top) and of a cell that is lowly blebbing (bottom) using optical tweezers. Right, plot displaying the relative trap force measured over time of the corresponding cells displayed on the left. Higher blebbing cells display higher variance of trap force over time compared to low blebbing cells.The scale bars represents 10μm. A red target has been placed at the initial position of the bead before tether formation to help visualize bead displacement. **(g)** The variance of the trap force over time and the number of blebs in a cell, indicating a strong correlation between the variance of membrane tension and blebbing.

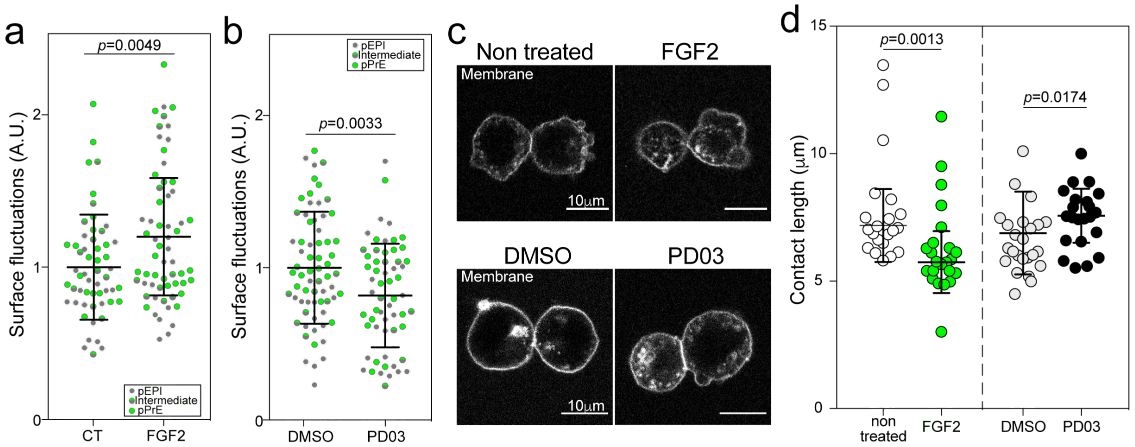


**Fig. S7 pPrE cells have higher FGF-mediated surface fluctuations than pEPI.**
**(a, b)** Single E3.75 pEPI and pPrE surface fluctuations with or without FGF2, 0.01% DMSO or PD03 treated for < 45 minutes. Each plot is a combination of N = 3 independent experimental results. The amplitude of surface fluctuations was calculated using images every ten seconds over a total of five minutes. The amplitude was normalised by the total mean of CT or DMSO surface fluctuations in each individual experiments. P-value calculated by 3-way ANOVA using cell type,treatment, and replicate number as variables, reported p-value is for treatment. **(c)** Representative images of E3.75 ICM cells doublets, treated with FGF2 or PD03, formed by dissociated single cells from E3.75 blastocysts. E3.75 ICM cells contain both pEPI and pPrE. Images were taken as stills from movies. Plasma membrane labelled with membrane dye, CellMask Deep Red (false-coloured white). **(d)** The contact length of E3.75 ICM cells doublets treated with or without FGF2, DMSO or PD03, treated for < 45 minutes. The contact length was calculated using images every one minute over ten minutes. Note that contact length was used here as opposed to angle because the FGF treated doublets were very blebby, which made it difficult to assess contact angle. Each plot is a combination of N = 4 independent experimental results. Each dot represents a doublet. P-value calculated by 2-way ANOVA using treatment and replicate number as variables, reported p-value is for treatment.

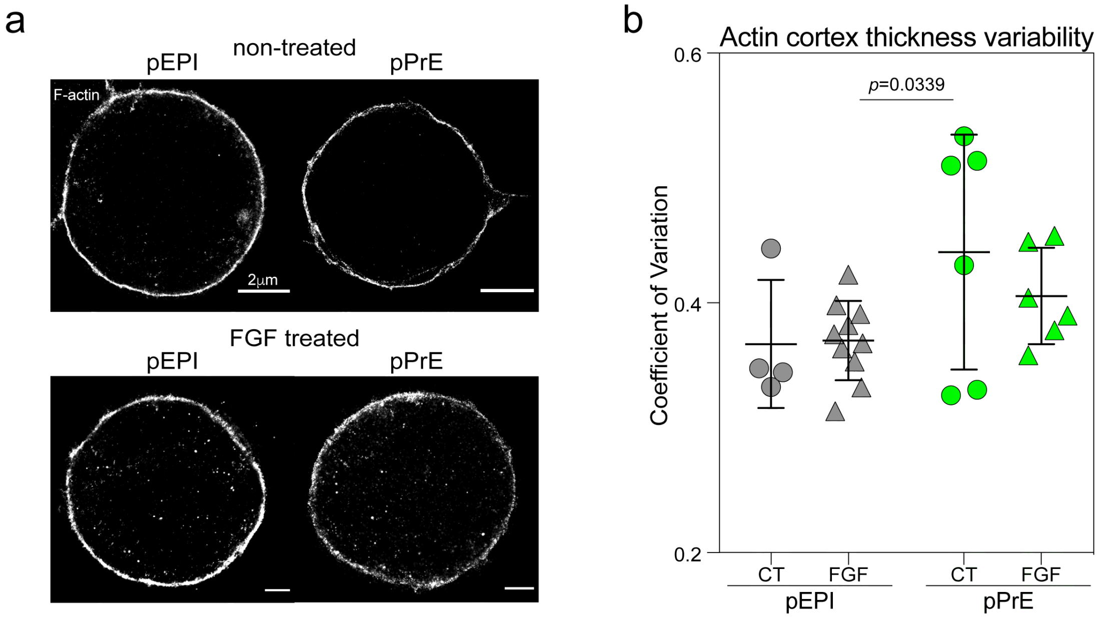


**Fig. S8: The actin cortex is more variable in pPrE cells.**
**(a)** Representative stochastic optical reconstruction microscopy (STORM) images of actin in *Pdgfra*$^{H2B\text{-}GFP/+}$ E3.75 pEPI and pPrE with and without FGF2. **(b)** Coefficient of variation of F-actin thickness in E3.75 pEPI and pPrE with and without FGF2 representing how variable the actin cortex was. Each dot represents a single cell. P-value was calculated by 2-way ANOVA using cell type and treatment as variables. Notably, there was no significant difference in variability when treating with FGF; however, the thickness of the cortex was significantly greater when the cells were treated with FGF (191 nm compared to 155 nm, $p \ll 10^{-4}$). We also note that the preparation for STORM requires many wash steps that may result in the cells with the most blebs being washed off.
-

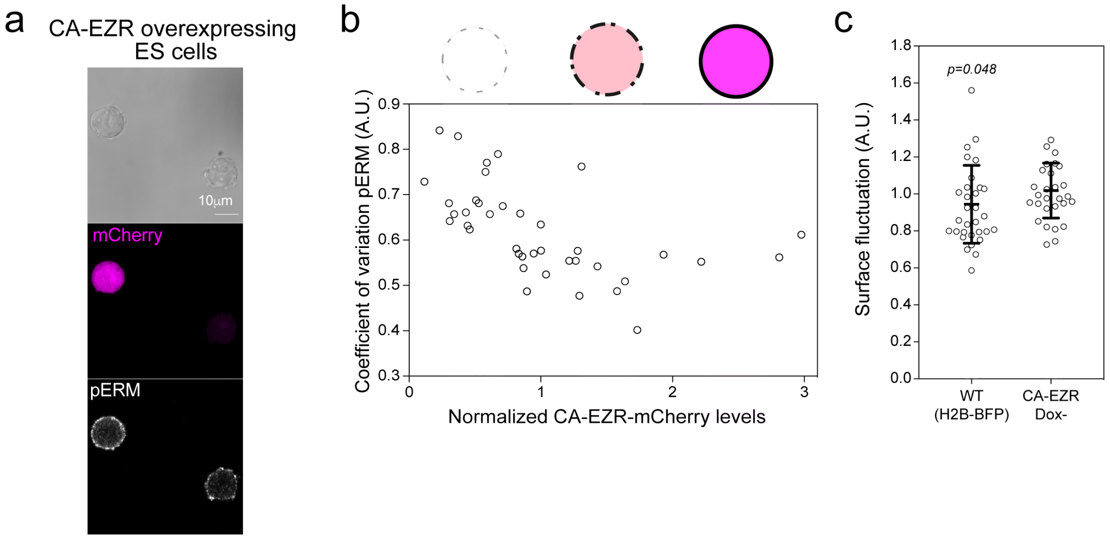


**Fig. S9: Engineered CA-EZR provides a range of surface fluctuations in ES cells.**
**(a)** Representative images of constitutively active Ezrin-IRES-mCherry (CA-EZR) ES cells, showing a high degree of pERM variability in the low mCherry expressing ES cells. **(b)** Schematic showing how pERM expression changes depending on CA-EZR expression. The coefficient of variation of pERM intensity along cell the surface is plotted against the intensity of mCherry. **(c)** Surface fluctuations of single cells of CA-EZR ES cells without Dox and H2B-BFP ES cells in 2i+LIF. P-value assessed by one-way ANOVA.

### Supplementary Materials and Methods

**Immunofluorescence staining**

Embryos and isolated ICMs and were fixed and stained in the same manner of embryos staining described in the Materials and Methods. After blocking, primary antibodies were diluted in blocking buffer and samples were incubated in the appropriate antibody solution at 4ºC overnight. They were rinsed three times in an embryo blocking buffer for 15 minutes ~ each. Secondary antibodies were diluted in blocking buffer with or without 500 ng/ml DAPI and samples were incubated in the appropriate antibody solution at room temperature for one hour in the dark. They were rinsed three times in an embryo blocking buffer for 15 minutes ~ each. Then the samples were transferred on PDL-coated glass-bottom dishes. Otherwise, the samples were incubated briefly in increasing concentrations of Vectashield before mounting on glass slides in small drops of concentrated Vectashield. Subsequently, coverslips with Vaseline spacer were mounted and sealed with nail varnish.

For pMRLC staining, cells were fixed and stained as a similar manner of pERM staining described in the Materials and Methods. PBS containing 1% BSA and 10% donkey serum was utilised for blocking buffer.

Primary and secondary antibodies and their concentrations were listed in Supplementary Data Table 3.

**ICM cells migration assay**

E3.75 *Pdgfra*$^{H2B\text{-}GFP/+}$ positive embryos obtained from intercrossing of *Pdgfra*$^{H2B\text{-}GFP/+}$ and CD-1 or F1 hybrid were used. Isolated single ICM cells in Blast medium containing 1:10000 CellMask Orange (Life Technologies) were loaded into the BSA coated polydimethylsiloxane (PDMS) confinement devices with a fixed roof height of 8 μm, 9 μm or 10 μm. Confinement devices were designed to restrict cells between two glass plates, trapping the cells in the *z*-direction but allowing free movement in *x*- and *y*-direction, as first demonstrated by (*1*). To adapt the device for the use with very small cell numbers, we modified confinement channels as described in (*2*) by replacing the channels with pillars to create a constant, well-defined roof hight. Live imaging of the cells were performed with a 6x silicon objective (UPLSAPO60XS, Olympus) on an inverted microscope (Olympus FV1200) equipped with a humidified chamber at 37°C and 5% $CO_2$ . The bright field, GFP, CellMask images of the cells were taken every one, two or three minutes for up to ten hours.

**Surface tension measurement**

**Cell preparation**: E3.75 *Pdgfra*$^{H2B\text{-}GFP/+}$ embryos obtained from intercrossing of *Pdgfra*$^{H2B\text{-}GFP/+}$ and F1 hybrids were used. Isolated ICM cells were transferred to single-drop of N2B27 in a glass-bottom dish (FluoroDish, World Precision Instruments) and incubated for 30 minutes at 37°C and 5% $CO_2$. 2 ml M2 in the presence of 0.01% CellMask Deep Red Plasma membrane Stain (Thermo Fisher Scientific) was added to the dish. The mean GFP intensity of *Pdgfra*$^{H2B\text{-}GFP}$ was used to classify ICM cells' lineage and cell cycle stage. Mitotic cells and dead cells were excluded from the analysis. Samples were measured for no longer than two hours.

**Experimental setup**: Tension measurements were performed using a JPK CellHesion (JPK Instruments) mounted on an IX81 inverted confocal microscope (Olympus). Tipless silicon cantilevers (ARROW-TL1Au-50) were chosen with a nominal spring constant of 0.03 N/m. Sensitivity was calibrated by acquiring a force curve on a glass coverslip. Spring constant was calibrated by the thermal noise fluctuation method. *Z*-length parameter and setpoint force were set at 30 μm and 10 nN, respectively. Constant height mode was selected. The measurement was carried on by lowering the tipless cantilever onto an empty area next to a target cell. Once the cantilever retracted (by roughly 30 μm), it was positioned above the target cell and run a compression for 200 seconds. During the constant height compression, the force acting on the cantilever was recorded. After initial force relaxation, the resulting force value was used to extract surface tension. A confocal stack was acquired using an Olympus UPLANSAPO ×60/1.35 NA oil immersion objective (Olympus).

**Analysis**: The calculation of cortex tension is based on (*3*). Briefly, neglecting the angle of the cantilever with respect to the dish (~8°) and assuming negligible adhesion between cell, dish and cantilever, the force balance at the contact point reads:

$$\mathrm{T} = \frac{\mathrm{F}\left(\frac{r_{\mathrm{mid}}^2}{r_{\mathrm{c}}^2} - 1\right)}{2\pi r_{\mathrm{mid}}} \qquad (1)$$

where $r_{\mathrm{mid}}$ is the radius of maximum cross-sectional area of the selected cell, $r_{\mathrm{c}}$ is the radius of contact area between cell and cantilever and $F$ is the resulting force exerted by the cell on the cantilever. To avoid errors due to direct measurement of $r_{\mathrm{c}}$, the contact radius was calculated using the following formula (2)(*4*):

$$A_{\mathrm{c}} = A_{\mathrm{mid}} - \left(\frac{\pi}{4}\right) \mathrm{h}_{\mathrm{cell}}^2 \qquad (2)$$

where $A_{\mathrm{c}}$ is the contact area between cell and cantilever, $A_{\mathrm{mid}}$ is the cell maximum cross-sectional area and $h_{cell}$ is the cell height. $h_{cell}$ was calculated as described (*4*) from the radius, force and cantilever height during compression. The cantilever height during compression was obtained by subtracting the cantilever height difference on glass and the cantilever height difference during cell compression. $h_{cell}$ values were confirmed by CellMask$^{TM}$ Deep Red membrane confocal stack reconstruction (corrected for optical aberration) (*5*-*7*).

**Membrane tension measurements by optical trap**

E3.75 *Pdgfra*$^{H2B\text{-}GFP/+}$ positive embryos obtained from intercrossing of *Pdgfra*$^{H2B\text{-}GFP/+}$ and CD-1 were used. Isolated single ICM cells were transferred in the M2 drops on a PDL-coated μ-Dish 35 mm ibidi dish. The dishes were exposed to plasma surface treatment (Pico, diener). 50 μg/ml PDL drop was put at the centre of the dish. The coated dishes were incubated for 30 minutes ~ one hour at room temperature. The PDL drop was washed with M2 medium for three times. The cells were kept for 15 minutes in a humidified incubator at 37°C and 5% $CO_2$. M2 medium concanavalin-A coated (50 μg/ml) carboxyl latex beads (1.9 μm diameter, Thermo Fisher Scientific [C37278]) were added to the drop prior to measurement.

A tether pulling assay was then performed using a homemade built optical tweezer (4W1064nm Laser Quantum Ventus) on an inverted microscope (Nikon Eclipse TE2000-U) equipped with a motorised stage (PRIOR Proscan). During the measurements, the position of the bead was recorded at a 90 milliseconds interval rate in the bright field using a 100x oil immersion objective (CFI Plan Fluor DLL, Nikon). pEPI and pPrE cells were identified by their GFP signals. The trap force was later calculated based on a product of the bead displacement (estimated using a custom-made ImageJ plugin) and of the trap stiffness (which was extracted using the method described in (*8*)).

For the measurement of the variance of the trap force overtime, ~$10^6$ ES cells were plated 16 hours prior to the measurement onto μ-Dish 35 mm ibidi dishes in N2B27. Membrane tension was then measured following the method described above. Of note, here membrane tension was continuously measured on the same cell using a single tether for approximatively five minutes. The number of blebs was then manually counted during the analysis, only blebs forming or retracting during the measurements were taken into account. Data in which the formation of blebs was physically interfering with the tether were excluded from the analysis.

**Surface fluctuation analysis of ES cells**

For CA-EZR-IRES-mCherry ES cells and H2B-BFP ES cells were plated for 24 hours in N2B27+2i+LIF media at $2 \times 10^4$ cells/cm$^2$, and the media were changed to N2B27+2i+LIF media. Then, cells were dissociated into a single-cell suspension using Accutase and suspended in 100 µl each culture media with 0.01% CellMask™ Green Plasma membrane Stain (Thermo Fisher Scientific). The media were buffered in the incubation chamber for at least 30 minutes before suspension. The analysis was performed as outlined in Supplementary information Fig.4.

**Analysis of doublet contact area**

E3.75 embryos obtained from intercrossing of CD1 were used for doublet formation. Doublets were made from isolated single ICM cells in the micro drop of Blast with or without 25 ng/ml FGF2, 1 µM PD03 or 0.01% DMSO and incubated for 30 minutes at 37°C and 5% $CO_2$. Doublet cells were transferred in Blast drop with or without 25 ng/ml FGF2, 1 µM PD03 or 0.01% DMSO in the presence of 0.01% CellMask™ Deep Red Plasma membrane Stain under the mineral oil in a PDL coated glass-bottom dish. Live images were acquired using a Leica TCS SP5 confocal microscope. A HC PL APO 40×/1.30 Oil CS2 with immersion oil was used. The middle section of the doublet images was taken every 20 seconds for ten minutes. The microscope is equipped with an incubation chamber to keep the sample at 37°C and 7% $CO_2$. The length of the cell-cell contact was measured every three frames (every one minute) for ten flames using Fiji, and was used as a proxy measurement of cell-cell contact area. The mean of measured contact length of a doublet was used for the plot (Fig. S7d).

**Direct stochastic optical reconstruction microscopy (dSTORM) imaging and analysis**

**Sample preparation:** E3.75 *Pdgfra*$^{H2B\text{-}GFP/+}$ positive embryos obtained from intercrossing of *Pdgfra*$^{H2B\text{-}GFP/+}$ and CD-1 were used. Isolated pPrE and pEPI cells were cultured in a PDL-coated small drop of N2B27 with or without 25 ng/ml FGF2 on 35 mm glass-bottom dishes (MatTek; P35G-0.170-14-C) at 37°C and 5% $CO_2$ for 45 minutes. The samples were fixed and stained in the same manner as pERM staining described in the Materials and Methods. Primary and secondary antibodies were diluted in PBST (PBS supplemented with 0.1% Tween). For the secondary antibody incubation, cells were first incubated with AlexaFluor donkey anti-goat 488 (Thermo fisher, 1:1000) followed by incubation with Hoechst (INVITROGEN, 1 µg/ml) and AlexaFluor647 Phalloidin (Thermo Fisher, 1:200) diluted in PBST for another hour. Cells were then gently washed three times with PBST. The imaging dishes were then filled with STORM buffer (50 mM Tris pH 7.5, 10 mM NaCl, 10% glucose (w/v), 27 mM MEA, 40 µg/mL catalase, 5 U/mL pyranose oxidase and 2 mM cyclooctatetraene) and imaged immediately.

**Imaging and analysis:** dSTORM imaging was performed on a commercial Zeiss Elyra 7 microscope. F-actin was imaged in the cellular mid-plane through a 63 x 1.46 NA alpha Plan-Apochromat oil-immersion objective lens and a 1x tube lens under 642 nm (100% laser power) and 405 nm (0-2% laser power) excitation. For each super-resolution image of F-actin, a Hoechst (405 nm excitation) and GFP (488 nm excitation) snapshot was also acquired. Fluorescence was captured on a sCMOS camera using 20 milliseconds integration time. A total of 15000 frames were typically acquired for super resolution image reconstruction. Reconstructions were generated using the in-built ZEN Black software (Zeiss). Briefly, single molecule localisation was estimated using a multiple object 2D Gaussian fitting routine, accounting for overlapping fluorophores. Data was post-processed to correct for mechanical drift during the acquisition by applying a model-based cross-correlation method and localisations with uncertainties >50 nm were then removed from the data-sets. Cortex thickness measurements were performed on the super-resolution images using a custom-written MATLAB script (described in (*9*)). Briefly, the cell cortex for each super-resolution data set was manually estimated using a custom MATLAB GUI. The cell cortex was then automatically detected by calculating the maximum intensity peak of the transverse intensity profile along the cell periphery at each pixel along with the user-supplied cortex co-ordinates. The cortex was then straightened using cubic spline fitting, and line scans of the intensity profiles across well-defined cortical regions were performed, from which full-width at half-maximum (FWHM) values were used to estimate cortical thickness.

##### Cell size measurement

EPI and PrE cells' diameters were measured using the images of the middle plane of the cells. For E3.5 cells, six EPI and six PrE cells were measured. For 3.75 cells, 21 pEPI and 22 pPrE cells were measured. For E4.5 cells, ten EPI and eight PrE cells were measured.

##### RNA extraction and cDNA synthesis

Total RNA was prepared with the RNeasy Kit (Qiagen), and reverse transcribed using SuperScriptII (Invitrogen) according to the manufacturer's protocol. Real-time PCR was performed using TaqMan Fast Universal Master Mix and TaqMan Gene Expression assays (Applied Biosystems). *Gapdh* was used as an endogenous control (Applied Biosystems). For siRNA experiments, the data were further normalised to control cell line. The TaqMan Gene Expression assays IDs were Mm00447761_m1; *Ezrin*, Mm00447889_m1; *Moesin*, Mm01177363_m1; *Radixin*.

**Statistical analysis**

For all statistical analysis in the paper, unless otherwise reported in corresponding figure legend, n-way ANOVA was used to calculate P-values to establish significant changes between any two means. Details of each ANOVA stated in figure legends: for example, variables used for the n-way ANOVA could be 'replicate number', 'cell type' (e.g. EPI and PrE) and when relevant 'small molecule treatment'. There was usually a significant interaction effect for 'experiment number' due to expected experimental variability in embryo work, so this $p$-value is suppressed throughout the paper and the $p$-value reported is for 'cell type' or 'small molecule treatment' depending on the experiment. The midline is mean of overall experiments, and error bars represent standard deviation over all experiments. For Fig. S7a, b, d, standard deviations were calculated as the square root of the sum of the squares of the standard deviation of each experiment divided by the number of experiments.

### Supporting movies legends

**Supplementary Data Movie 1:**

**EPI and PrE sorting in isolated E3.75 ICM.**

Time-lapse sequence of isolated E3.75 mTmG$^{+/-}$*Pdgfra*$^{H2B\text{-}GFP/+}$ ICM in an embryo immobilisation chip.

**Supplementary Data Movie 2:**

**Surface fluctuation of isolated E3.75 pEPI.**

Time-lapse sequence of isolated E3.75 mTmG$^{+/-}$*Pdgfra*$^{H2B\text{-}GFP/+}$ pEPI on PDL-coated dish. This file was assembled at the middle section of the cell every ten seconds over five minutes.

**Supplementary Data Movie 3:**

**Surface fluctuation of isolated E3.75 pPrE.**

Time-lapse sequence of isolated E3.75 mTmG$^{+/-}$*Pdgfra*$^{H2B\text{-}GFP/+}$ pPrE on a PDL-coated dish. This file was assembled at the middle section of the cell every ten seconds over five minutes.

**Supplementary Data Movie 4:**

**Surface fluctuation of E3.75 ICM aggregates.**

Time-lapse sequence of isolated E3.75 mTmG$^{+/-}$*Pdgfra*$^{H2B\text{-}GFP/+}$ ICM aggregates. This file was assembled at one Z-section of the aggregation of every 20 seconds over five minutes. (a) ICM aggregate 1. (b) ICM aggregate 2.

**Supplementary Data Movie 5:**

**Surface fluctuation of isolated E3.75 pEPI with FGF2.**

Time-lapse sequence of isolated E3.75 mTmG$^{+/-}$*Pdgfra*$^{H2B\text{-}GFP/+}$ pEPI with FGF2 on PDL-coated dish. This file was assembled at the middle section of the cell every ten seconds over five minutes.

**Supplementary Data Movie 6:**

**Surface fluctuation of isolated E3.75 pPrE with FGF2.**

Time-lapse sequence of isolated E3.75 mTmG$^{+/-}$*Pdgfra*$^{H2B\text{-}GFP/+}$ pPrE with FGF2 on a PDL-coated dish. This file was assembled at the middle section of the cell every ten seconds over five minutes.

### Supplementary Data Tables legends

**Supplementary Data Table 1:**

**(a)** Actin-cytoskeleton genes list based on Go terms. **(b)** Actin-cytoskeleton genes based on the listed Go terms.

**Supplementary Data Table 2:**

Highly modulated actin-cytoskeleton related genes in ICM cells through E3.5 to E4.5

**Supplementary Data Table 3:**

Antibody list.

**Supplementary Data Table 4:**

ERM siRNA or NC siRNA treated ES cells location and numbers in the chimaera blastocyst.

### Extended Supplementary Information

##### Doublet formation from isolated ICM cells

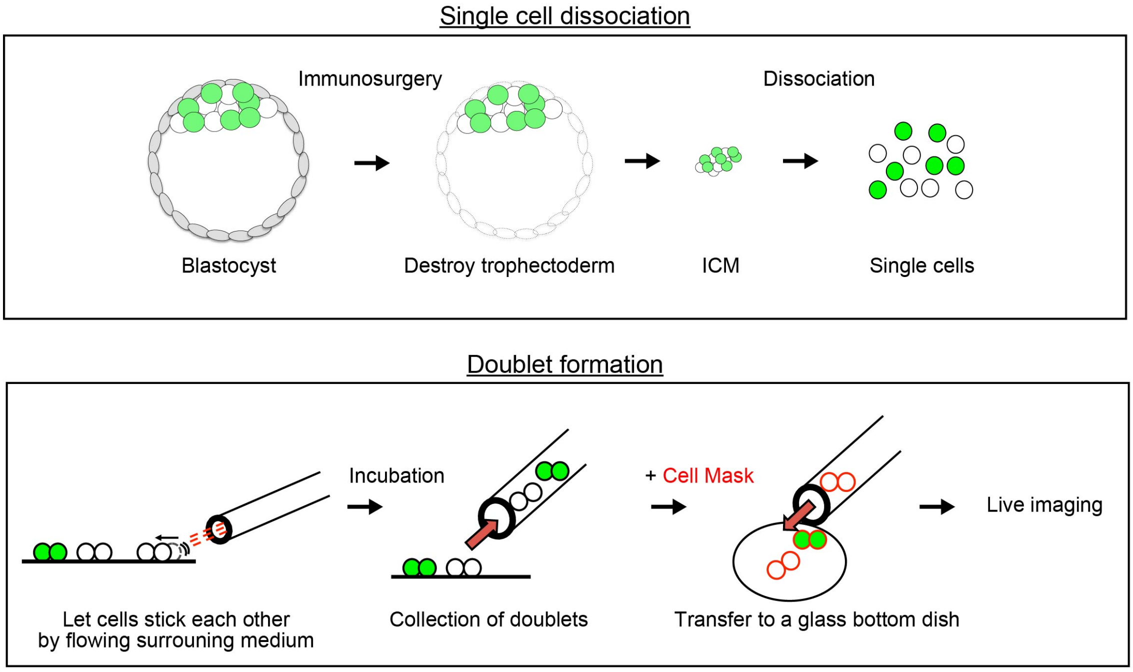


**Extended Supplementary information Fig. 1: Schematic images of pEPI and pPrE cells doublets formation.**

##### CS3D method: modelling surface fluctuations and blebbing

Simulation results in this work were produced by the model presented in reference (*10*) which we call cell sorting in 3D (CS3D, http://github.com/chris-revell/SEM). CS3D is an extension of the Subcellular Element Method (*11*), which is a force-based technique for modelling the development of multicellular tissues. It allows us to study the effects of complex inter- and intra-cell features on tissue-scale dynamics (*12*). Each individual cell is modelled as a group of infinitesimal elements, interacting via nearest-neighbour forces (Extended Supplementary information Fig.2a, b) defined by Morse potentials (*13*). Nearest-neighbour elements of different cells interact by the same mechanism. This model produces a relatively fine-grained representation of multicellular systems incorporating both inter- and intra-cellular mechanisms.

For CS3D, we implemented a simple algorithm for identifying boundary elements in each cell and used a Delaunay triangulation (*14*, *15*) over this set of elements to define a nearest neighbour network across the cell boundary. By applying a constant force between these neighbouring elements in the triangulation, we modelled cell cortical tension (Extended

Supplementary information Fig.2c). The magnitude of tension forces defined within the Delaunay triangulation can vary locally across the cell surface at interfaces with surfaces of other cells. We define a cell-cell interface by identifying and labelling the cortex elements that share adhesive inter-cell interactions with cortex elements of another cell.

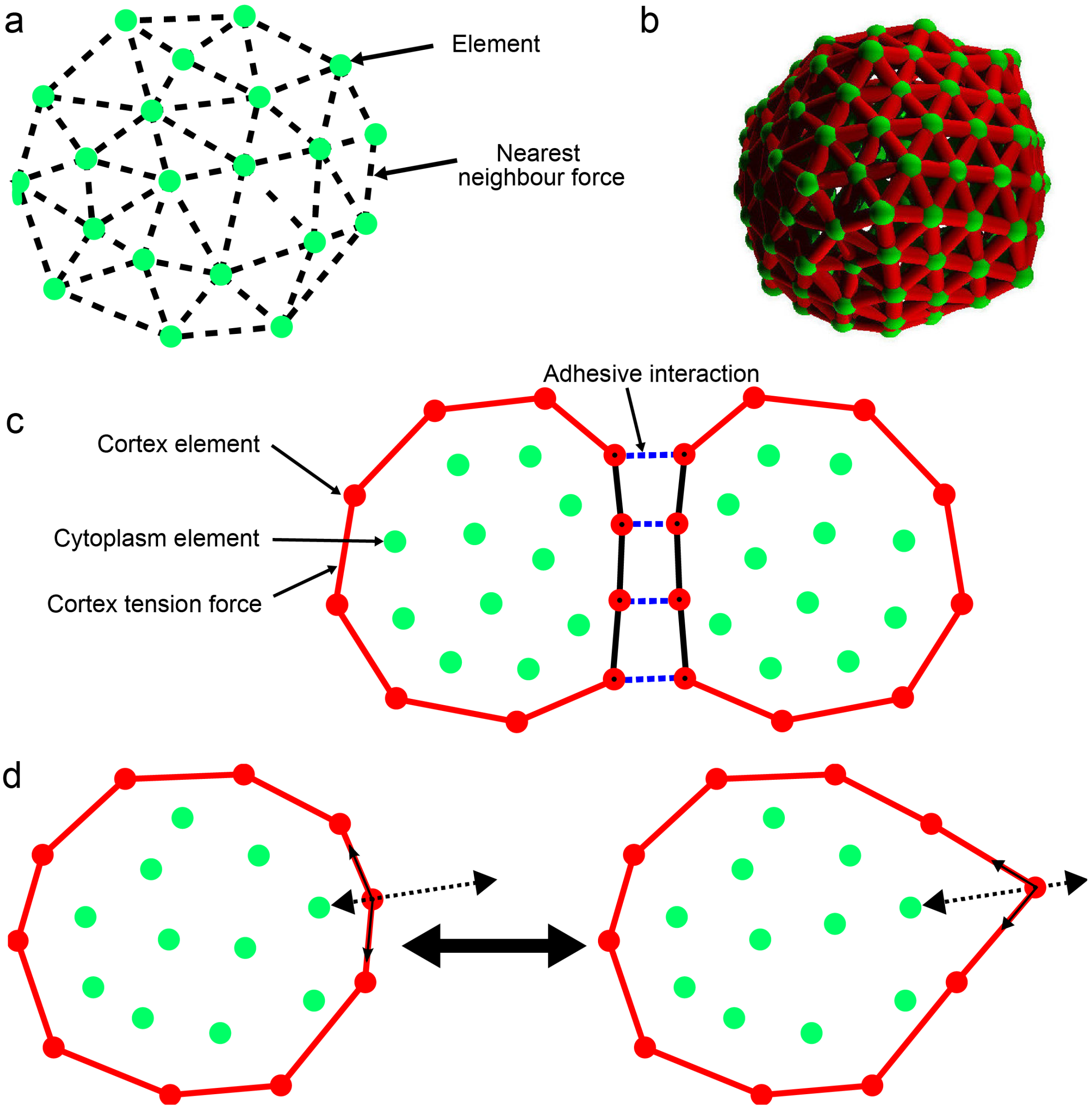


**Supplementary information Figure2.**
**(a)** Schematic of cell in 3D force-based cell sorting simulation (CS3D) method, showing elements as green circles, and highlighting local interactions between elements with dotted lines. **(b)** Visualisation of one cell showing boundary elements to define contractile cortex. **(c)** Schematic demonstrating intercellular interactions as laid out[10]. Intercellular adhesive interactions (blue) define interface, and tension is defined for cortex (red). **(d)** Implementation of surface fluctuations in PrE. Cortical tension is varied sinusoidally in time, causing the element to protrude from the cell surface before being retracted.

Importantly, in reference (*10*), we show that there is a nearly perfect correlation between the contact area (i.e. the affinity) and the level of sorting in a multicellular aggregate, regardless of the underlying force balance that gives rise to that contact area. Therefore, based on this simplification and the results of Fig. S4a, both cell types were given the same cortical tension and the same adhesion magnitude (approximated as $0.2\gamma_c$ as inspired by references (*16*, *17*), where $\gamma_c$ is the cortical tension), and the only parameter that was varied was interfacial tension. For simplicity, all interfacial tensions were set equal to unity other than EPI::EPI interfacial tension, which determines the affinity parameter. All tests we ran indicated that the only value that played a role in sorting was the ratio between the interfacial tensions of EPI::EPI and PrE::PrE and not the absolute values. The median values of pEPI and pPrE cell's external contact angle are used to estimate the dimensionless parameter $\beta$ given by equation (1) in the text. The value of $\cos(\theta_{\mathrm{EPI}})$ was ~0.59, and the value of $(\theta_{\mathrm{PrE}})$ was ~0.79 leading to $\beta \approx 0.75$.

We model surface fluctuations, $\delta$, as a local change in the cortical tension resulting in a protrusion from the cell surface. To achieve this, we devised a simple algorithm demonstrated in Supplementary information Fig.2d. Each cortex element on the surface of a cell is given a randomly allocated phase $\phi$, which increases linearly with time. This phase is used to determine the strength of cortex forces experienced by the cortex element, which vary sinusoidally with a period of $\tau/10$ where $\tau$ is the cell cycle time. Thus, the force experienced by the element from any cortical tension interaction is modulated by a term, $\delta * \sin(10t/\tau + \phi)$. A dimensionless parameter $\varepsilon$ can be formed from the ratios of $\delta$ in pPrE to pEPI. This oscillation in tension at a particular point in the surface causes the element to protrude from the cell surface before being pulled back in, modelling a fluctuation. The behaviour of the system can be controlled by varying the surface fluctuation amplitude. The relevant parameter is the ratio of surface fluctuations of PrE to EPI, $\varepsilon$, which we calculated as $\approx 0.35$ ($\delta_{\mathrm{EPI}} = 1.39$ μm, $\delta_{\mathrm{PrE}} = 1.91$ μm).

Throughout each simulation, the extent of sorting was probed using a numerical sorting index (*10*). The sorting index is the ratio of the proportion of this area occupied by PrE cells to the total external surface area of the cell aggregate (Extended Supplementary Information Fig.3a). To provide context to the values obtained from the surface sorting measure, we implemented a randomised system normalisation (*10*). This algorithm randomly reallocates the fates of all cells in the system after each measurement, retaining the spatial arrangement and the number of each cell type, and repeats the measurement for each arrangement 100,000 times to obtain the mean and standard deviation of the measure across all randomised systems (Extended Supplementary Information Fig.3b). The system is then reverted to the original simulation state. We can then present the results of the simulation as *Sorting index* = $\frac{X_S-\mu}{4\sigma}$. Each parameterisation of the model was tested four times for all results in the paper, and the mean and standard deviation for each run is reported.

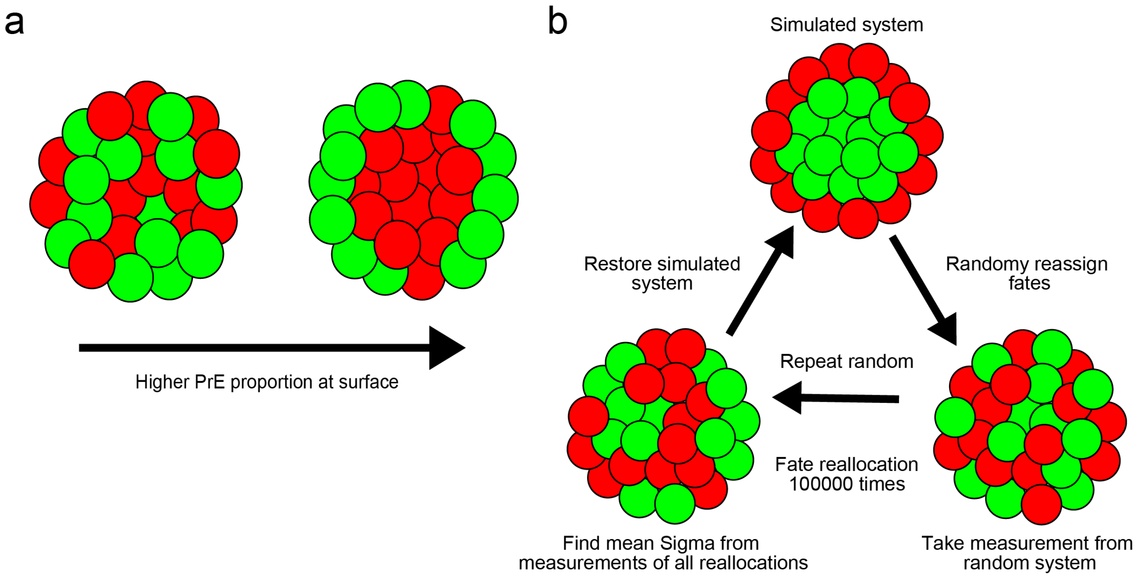


**Extended Supplementary information Fig. 3**
**(a)** Diagram demonstrating change in surface occupancy of PrE (green) in sorted systems. The proportion of the external system surface occupied by PrE in calculated for the surface sorting measure. **(b)** Diagram demonstration how fates of cells are randomised before repeating measurements in order to find mean and standard deviation across random systems, which are then used to calculate sorting indices.

In the simulation, we set pEPI and pPrE cells size (radius) as the same based on the measurement (E3.5 pEPI/ICM: 7.22±0.69, pPrE: 7.23±1.22, E3.75 pEPI: 7.13±1.31, pPrE: 7.27±1.19, E4.0 pEPI/EPI: 6.88±0.66, pPrE/PrE: 6.26±0.83). EPI and PrE in both a half embryo and a double embryo can sort similarly to a normal size embryo indicating that the total number of ICM cells, at least from 0.5 to 2 times difference, doesn't affect EPI and PrE segregation (*18*). We simulated up to 50 cells noting that the number of E3.5 ICM and E4.5 ICM is roughly 10-20 cells and 40-50 cells respectively. We approximated the proportion of both pEPI and pPrE as 50%.

Simulations were performed on the University of Cambridge Darwin HPC facility, running one simulation per core independently, using the Intel FORTRAN compiler.

**Quantification of membrane dynamics**

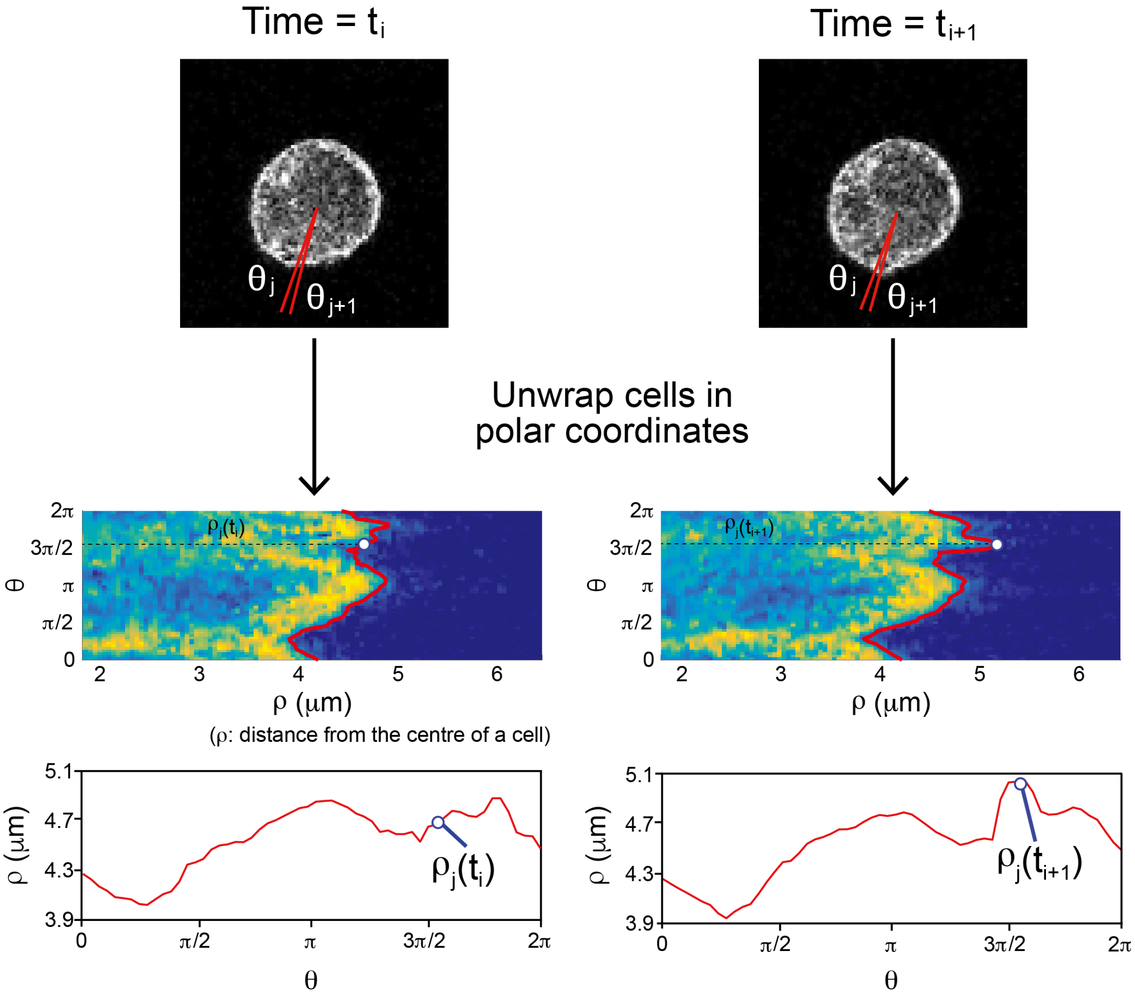


$$V_j = (SD(\{\rho_j(t_1), \rho_j(t_2), \ldots, \rho_j(t_N)\}) \quad j = 1{:}M$$

$$\text{Surface fluctuations} = \text{Mean}(\{V_1, \ldots V_M\})$$

**Extended Supplementary information Figure. 4:**
**The analysis method of membrane dynamics of isolated pEPI and pPrE cells.**
Schematic outline of how surface fluctuation of a cell is measured from cell imaging data.

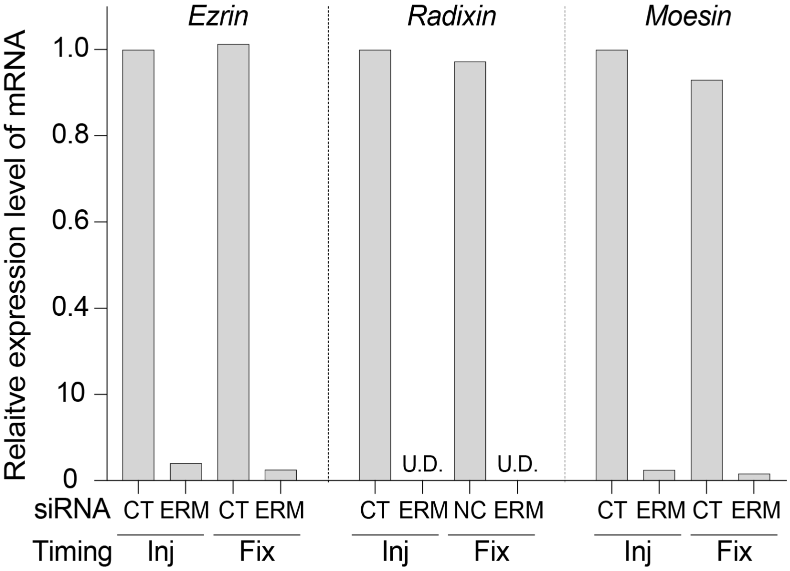


**Extended Supplementary information Fig. 5: Ezrin, Radxin and moesin knockdown efficiency**
The mRNA expression levels of *Ezrin*, *Radxin* and *Moesin* in Negative control siRNA mix treated ESCs (N) and Ezrin, Radxin and moesin siRNAs treated ESCs (ERM). mRNA expressing levels were normalised using the expression of *gapdh*. U.D.: undetectable level. Inj: the siRNA treated ESCs were collected at the same timing of the blastocyst injection. Fix: the siRNA treated ESCs were collected at the same timing of the chimaera embryo fixation. **(j)** pMLRC expression in wild-type ESCs and CA-EZR ESCs.

### Extended Supplementary Information References

#### Abbreviations

AFM : atomic force microscopy

ANOVA : analysis of variance

A.U. : arbitrary unit

BFP : blue fluorescent protein

BSA : bovine serum albumin

CA-EZR : constitutive active form of Ezrin

Cdx2 : caudal-related homeobox transcription factor 2

CS3D : 3D force-based cell sorting simulation

CSB : cytoskeletal stabilizing buffer

CV : coefficient of variation

DAPI : 4',6-diamidino-2-phenylindole

Dim : dimension

DMEM/F-12 : Dulbecco's Modified Eagle's Medium/Nutrient Mixture F-12 Ham

DMSO : Dimethyl sulfoxide

Dox : doxycycline

E3.75 : embryonic day 3.75

EB : embryoid body

EPI : epiblast

ES cell : embryonic stem cell

F-actin : filamentous actin

F1 : first filial generation

FBS : foetal calf serum

FGF : fibroblast growth factor

FPKM : fragments per kilobase of exon per million mapped fragments

Gata : GATA-binding factor

GFP : green fluorescent protein

GMEM : Glasgow's minimum essential medium

GO term : Gene Ontology term

h : hour

H2B : human histone H2B

ICM : inner cell mass

IRES : internal ribosome entry site

LIF : leukaemia inhibitory factor

mTmG : membrane-localised tdTomato membrane-localised GFP

Nanong : nanog homeobox

NEAA : non-essential amino acids

Neo : neomycin

pERM : phospho-Ezrin/Radixin/Moesin

PBS : phosphate buffered saline

PCA : principal component analysis

PDL : Poly-D-lysine

PD03 : PD0325901, MEK inhibitor

Pdgfra : Platelet-derived growth factor receptor alpha

pEPI : EPI-precursor

pERK : phosphorylation of Extracellular signal-related kinase

pERM : phosphorylation of ezrin, radixin, moesin

PFA : paraformaldehyde

pPrE : PrE-precursor

PrE : primitive endoderm

PVP : polyvinylpyrolidone

RNA : ribosomal ribonucleic acid

RNA-seq : RNA sequencing

RT-qPCR : real time-quantitative polymerase chain reaction

Sox : SRY (sex determining region Y)-box

tdTomato : tandem dimeric Tomato
